# Targeted lipid A modification of *Shigella* vaccine strains reduced endotoxicity without compromising immunogenicity or invasiveness

**DOI:** 10.1101/2023.08.24.554604

**Authors:** Matthew E Sherman, Jane Michalski, Sayan Das, Hyojik Yang, Shoshana Barnoy, Lakshmi Chandrasekaran, Malabi Venkatesan, Robert K Ernst

## Abstract

*Shigella* infection contributes significantly to the global disease burden, especially affecting young children in developing countries. Currently, a vaccine against *Shigella* is unavailable and the prevalence of antibiotic resistance amongst *Shigella* species is continually rising. Live-attenuated *Shigella* vaccine candidates developed at Walter Reed Army Institute of Research have shown remarkable immunogenicity but exhibit adverse reactogenicity, most likely due to the highly toxic lipid A moiety present on the bacterial membrane. Previous attempts at reducing the endotoxicity have focused on deletion of intrinsic lipid A biosynthesis enzymes. In this study, we instead introduce exogenous lipid A modifying enzymes, generating targeted modifications in the lipid A structure, leading to a dampened TLR4 response within the host. In doing so, we generated vaccine candidates with detoxified lipid A and unaltered O-antigen structure thereby preserving the serotype-specific immunity while reducing endotoxicity.

## Introduction

*Shigella* are Gram-negative bacteria known to cause diarrhea and dysentery in humans through ingestion of contaminated food and water^1–4^. In children under 5 years of age, the uncontrolled inflammation and severe dehydration can lead to growth abnormalities, seizure, and even death^2,5–8^. The clinical severity^9,10^ and emergence of antibiotic resistance^11,12^ has prompted the development of multiple *Shigella* vaccine candidates that are currently in preclinical and clinical phases^4,13^; however, to date there is no licensed *Shigella* vaccine. Ongoing vaccine strategies for shigellosis include glycoconjugate vaccines, mainly composed of the O-antigen portion of lipopolysaccharide (LPS) conjugated to carrier proteins (or adjuvants)^4,13–16^ or live-attenuated and inactivated oral bacterial vaccines^4,13–15,17^ where the O-antigen is presented in its native context on the outer membrane of the bacterial cell. Protection against shigellosis is largely serotype-specific^4,13–15^ implicating the O-antigen as being the critical antigenic target for vaccine development. Therefore, vaccine candidates have focused primarily on *S. flexneri* 2a, 3a, and 6 as well *S. sonnei,* which are the predominant disease-causing serotypes^18^ and whose O-antigen structures are well characterized^14^.

Live-attenuated vaccines have the advantage of mimicking a natural *Shigella* infection and have been previously developed through serial passage or incorporation of targeted genetic mutations^13,15^. Both methods have generated vaccines that induce protective immune responses against virulent challenge in volunteer studies^19^; however, significant issues such as adverse reactogenicity has slowed the progress towards a universally accepted safe and effective *Shigella* vaccine^15^. Live-attenuated vaccine candidates have advanced into clinical trials by multiple academic, industrial, and government laboratories^4,13,15^. For this study, Walter Reed Army Institute of Research (WRAIR) has constructed several iterations of live-attenuated *Shigella* vaccines with targeted deletions in virulence-associated genes^20–28^. During clinical trials, these vaccines induced protective immunity in adults and children, but reactogenic outcomes were observed at the moderate to high doses required to confer protective immunity^29–36^. To reduce these side effects, we chose to target the highly immunostimulatory LPS molecule present on the bacterial membrane, which is thought to be a major contributor to these adverse effects.

LPS is a glycolipid composed of three regions, the O-antigen, core oligosaccharide, and lipid A membrane anchor; however, the lipid A region is the immunostimulatory part of the molecule^37,38^. Through a series of accessory proteins, lipid A binds to the TLR4/MD-2 receptor complex, initiating downstream signaling, such as the NF-*κ*B pathway, ultimately driving pro-inflammatory cytokine production^39,40^. The TLR4/MD-2 response is primarily driven by the structural features of lipid A, which vary across Gram-negative bacteria^39,41–43^. *Shigella* contains the prototypical lipid A structure comprised of six acyl chains (hexa-acylated) connected to a di-glucosamine backbone flanked by two terminal phosphates (bis-phosphorylated)^43^. This structure is known to be a potent stimulator of TLR4/MD-2^44^ and the ensuing inflammation likely contributes to the febrile symptoms observed when volunteers orally ingest *Shigella* for vaccination. Thus, the immunostimulatory lipid A moiety was the primary target for detoxification, particularly since the live-attenuated *Shigella* vaccines used in this study already lacked their capacity to invade adjacent intestinal epithelia^21^ and were deficient for all known enterotoxins^23,26^.

In this study we employed bacterial enzymatic combinatorial chemistry (BECC)^45^ whereby lipid A modifying enzymes were ectopically expressed within both *Shigella* WT and vaccine strains generating custom-designed lipid A structures on the outer membrane that lacked acyl chains and/or terminal phosphates. Successful lipid A deacylation and dephosphorylation upon expression of BECC constructs was confirmed by MALDI-TOF MS and found to be extremely robust and stable. Ultimately, dephosphorylation of *Shigella* lipid A, rather than deacylation, reduced the potency of LPS-induced immune signaling and severely decreased the endotoxicity *in vivo*. Additionally, dephosphorylation of the lipid A moiety did not impair the invasion of colonic epithelia or immunogenicity in a mouse pulmonary model. Altogether, we reveal new live-attenuated *Shigella* vaccines strains, altered only in their lipid A region, which are greatly detoxified without any consequence to phenotypic traits.

## Results

### Development of Targeted Lipid A modifications in *Shigella* strains

We first tested the expression of BECC constructs *in trans* from the plasmid pSEC10M (Figure S1A, B) in wild-type *Shigella* and first-generation vaccine constructs WRSs1 (*S. sonnei*) and SC602 (*S. flexneri* 2a). The lipid A biosynthetic enzymes LpxE (phosphatase) and PagL (deacylase) were expressed alone and in combination (termed “Dual”) from the *ompC* promoter (P*_ompC_*) in wild-type and attenuated strains of *Shigella.* The expected lipid A modifications (Figure 1) were confirmed using FLAT^46^ processing and subsequent MALDI-TOF MS analysis (Figure S2). The expected lipid A phenotype for unmodified *Shigella* was confirmed to be a hexa-acylated bis-phosphorylated lipid A molecule with a base peak at *m/z* 1797. Expression of LpxE and PagL generated base peaks at *m/z* 1717 and 1570 representing lipid A structures lacking a phosphate or 3OH C14, respectively. The expression of Dual generated a minor peak at *m/z* 1490 indicative of a loss of both a phosphate and 3OH C14 acyl chain (Figure 1, S2). These results confirmed functional lipid A modification was achievable in *Shigella*.

**Figure 1:**
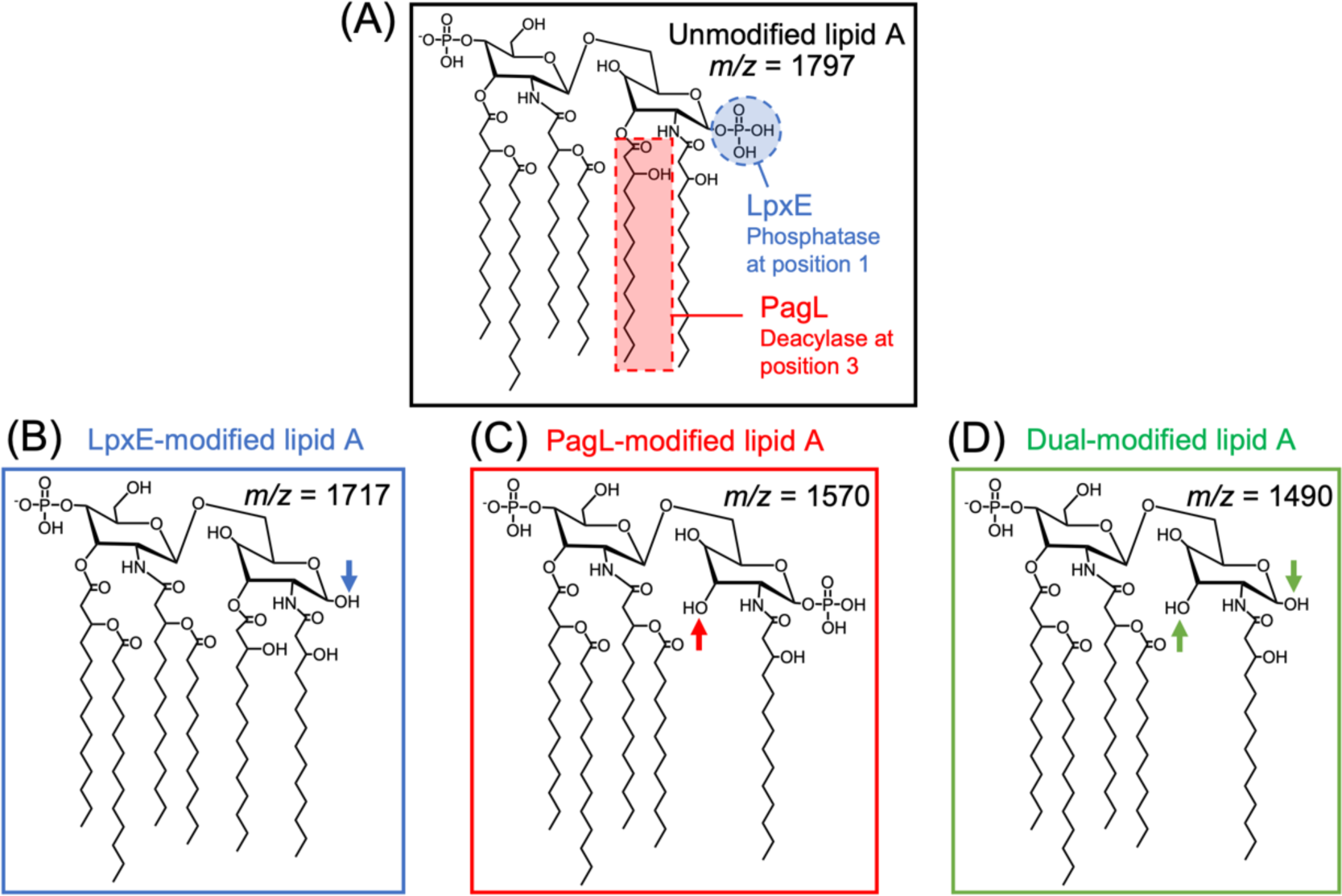
Lipid A modifications generated by BECC enzymes used in this study. (A) The native lipid A structure of *Shigella* along with the (B) LpxE-(C) PagL- and (D) Dual-modified resultant structures, lacking a phosphate, 3OH C14 acyl chain, or both, respectively. The expected *m/z* for the [M-H]^−^ ions observed in MALDI-TOF spectra is displayed for each structure.

We next integrated the BECC constructs into the *Shigella* chromosome to establish stability without the need for antibiotic selective pressure. Single-copy chromosome insertions were generated via Tn7 transposition into the *attTn7* site downstream from the *glmS* gene (Figure S1C) and confirmed by PCR and whole genomic sequencing. Integration of P*_ompC_*-*lpxE* and P*_ompC_*-Dual was achieved in both wild-type *S. sonnei* Moseley and *S. flexneri* 2a 2457T but integration into second-generation attenuated vaccine strains WRSs2 (*S. sonnei*) and WRSfG12 (*S. flexneri* 2a) varied. We were able to chromosomally integrate P*_ompC_*-*lpxE* into both WRSs2 and WRSfG12 but integration of P*_ompC_*-Dual was not achieved and deemed a lethal event to the already genetically manipulated *Shigella* strains. Chromosomal expression of BECC enzymes generated a pronounced enhancement of lipid A modification as complete shifts of the base peak from *m/z* 1797 to 1717 and 1490 were observed upon *lpxE* and Dual expression, respectively (Figure 2). To analyze any off-target effects of BECC modification, MALDI-TOF MS analysis in the range of *m/z* 500 – 1000 of total lipid extracts revealed no gross changes to the membrane lipid composition of any of the BECC-modified variants (data not shown).

**Figure 2:**
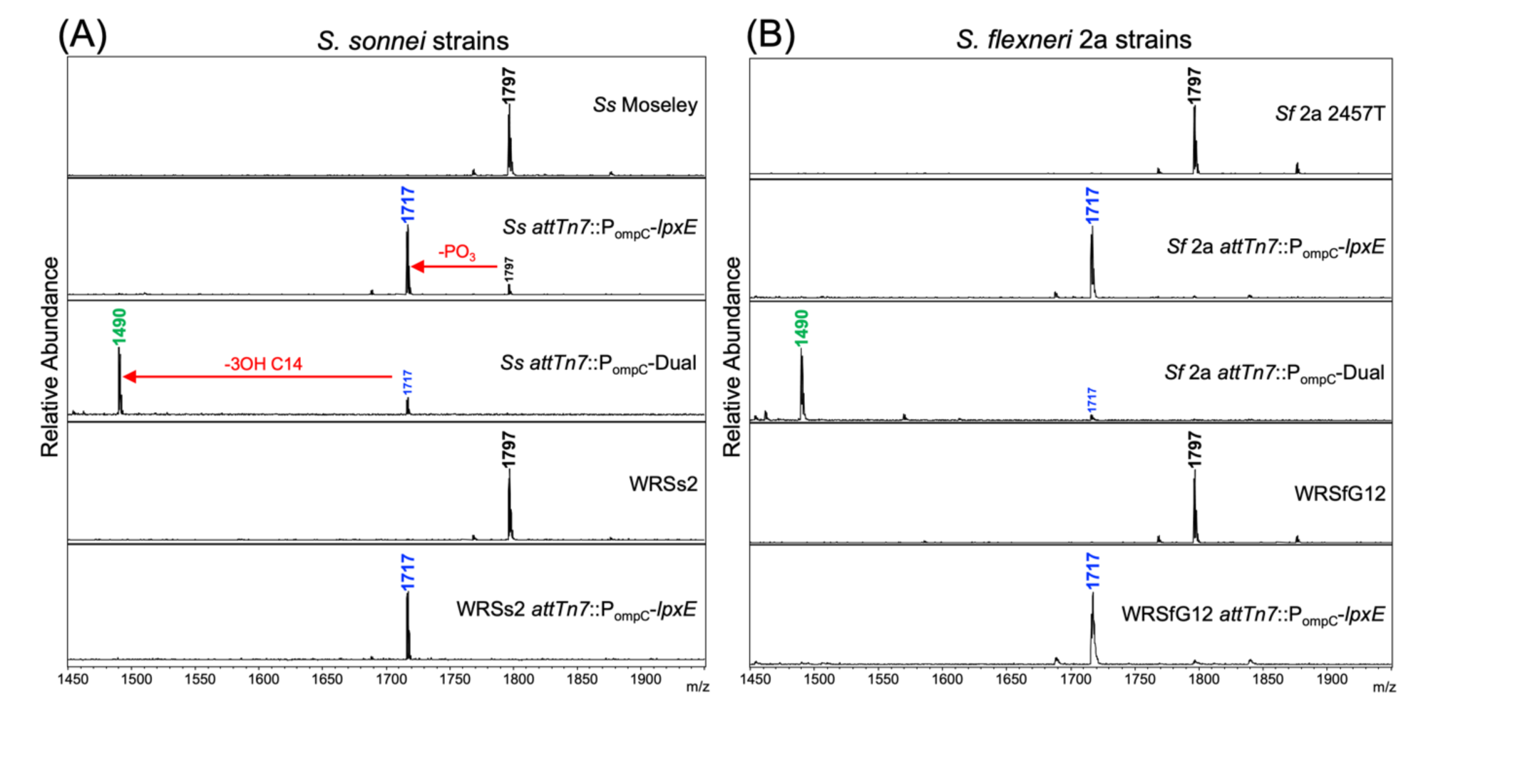
MALDI-TOF MS spectra of lipid A related peaks from BECC-modified *Shigella* strains. MALDI-TOF MS spectra using the FLAT method on (A) *S. sonnei* and (B) *S. flexneri* 2a wild-type and vaccine strains that were chromosomally integrated with BECC-constructs. Arrows depicted in (A) indicate the loss of a phosphate (PO_3_) or acyl chain (3OH C14) from the native lipid A structure.

### Site-specific Lipid A modifications confirmed using MS/MS

To determine site-specificity of the lipid A modifications, FLAT processing and subsequent MALDI-TOF MS/MS analysis (termed “FLAT^n^”)^47^ was performed on precursor ions *m/z* 1797, 1717, and 1490 for unmodified, LpxE-, and Dual-modified lipid A, respectively (Figure 3A). Fragmentation of these precursor ions and subsequent mass scanning from *m/z* 500 - 1900 resulted in several fragment ions (Figure 3B) that correlated by mass to fragmentation patterns from the lipid A precursor ions (Figure 3C). Fragmentation of the *m/z* 1797 precursor ion generated two signature ions; the B2 ion at *m/z* 1698.24 confirmed hexa-acylation, whereas the 3α ion at *m/z* 1552.01 confirmed the bis-phosphorylation of the lipid A. Fragmentation of *m/z* 1717 precursor ion generated a signature ion at *m/z* 690.40 that confirmed dephosphorylation at the 1-position, as opposed to the 4’-position. Fragmentation of *m/z* 1490 precursor ion generated a signature ion of *m/z* 750.42 that confirmed loss of a phosphate at the 1-position and loss of a 3OH C14 at the 3-position of *Shigella* lipid A (Figure 3D). Data shown in Figure 3 are representative spectra collected from BECC-modified Moseley; however, the data was replicated using BECC-modified 2457T as well (data not shown). Altogether, the MS/MS data confirmed that lipid A modifications had occurred at the predicted sites.

**Figure 3:**
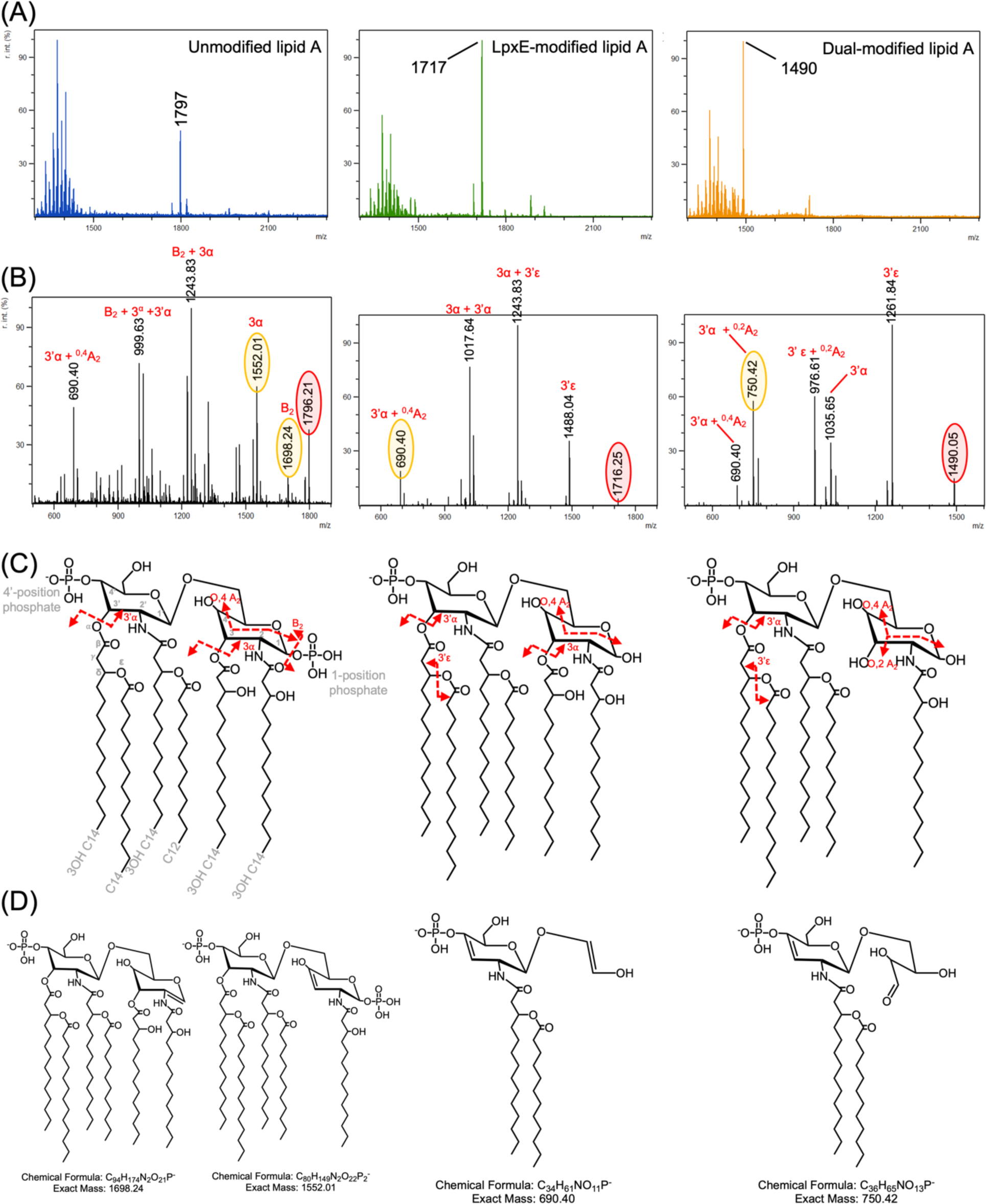
MS/MS analysis of unmodified and BECC-modified lipid A from *Shigella* strains. Lipid A variants were processed via the FLAT method and analyzed via MS/MS on a Bruker timsTOF. (A) Precursor ions, *m/z* 1797, 1717, and 1490, specific to unmodified, LpxE-, and Dual-modified lipid A molecules, respectively, were selected for fragmentation. (B) Resultant fragmentation of the precursor ions. Precursor ion and signature fragment ions are circled in red and yellow, respectively. Assigned fragment ions are shown in red above spectra (C) Lipid A structure showing predicted fragment ion breakages correlated to assignments. (D) Structures of the signature ions that resolve the site-specific modifications.

### Minimal effect of lipid A modifications on phenotypic traits of *Shigella*

To determine if there were any adverse phenotypic changes due to the BECC modifications, *Shigella* were assayed for bacterial cell growth and epithelial cell invasion. BECC-modified *Shigella* strains showed little to no altered bacterial growth compared to their unmodified counterparts over the course of 15 hours. All strains reached logarithmic growth within 5 hours and entered stationary phase within 6 hours (Figure S3). Cell density after 15 hours was not as high for WRSs2 *attTn7*::*lpxE* compared to its parental strain, WRSs2, but this did not affect LPS yields from cultures (data not shown).

Bacterial invasiveness was assessed using an established gentamicin protection assay^23,25,26^. At a multiplicity of infection (MOI) of 10, we found similar invasion patterns when comparing parental strains *S. sonnei* Moseley and WRSs2 to their BECC-modified counterparts. However, WRSs2 exhibited significantly (p ≤ 0.001) higher invasion (0.51 – 0.52%) than wild-type Moseley (0.08-0.11%) (Figure 4A). There was no significant difference in invasion between any of the *S. flexneri* 2a strains with the exception of Dual-modified 2457T (Figure 4B). *S. flexneri* 2457T *attTn*7::*P_ompC_-*Dual had significantly (p ≤ 0.05) lower invasion (0.41%) than unmodified 2457T (2.94%) due to a high frequency of invasion plasmid loss (data not shown). Assessment of IL-8 secretion in the cell culture supernatants followed the same pattern observed in the invasion assay. Significantly (p ≤ 0.05) higher IL-8 secretion levels were observed with invasion of WRSs2 compared to wild-type Moseley, but these levels were not impacted by BECC-modification (Figure 4C). Moreover, the reduced invasion observed with *S. flexneri* 2457T *attTn*7::*P_ompC_-*Dual resulted in decreased (39-fold) IL-8 secretion compared to wild-type 2547T (Figure 4D). These data were recapitulated at an MOI=100 (Figure S4).

**Figure 4:**
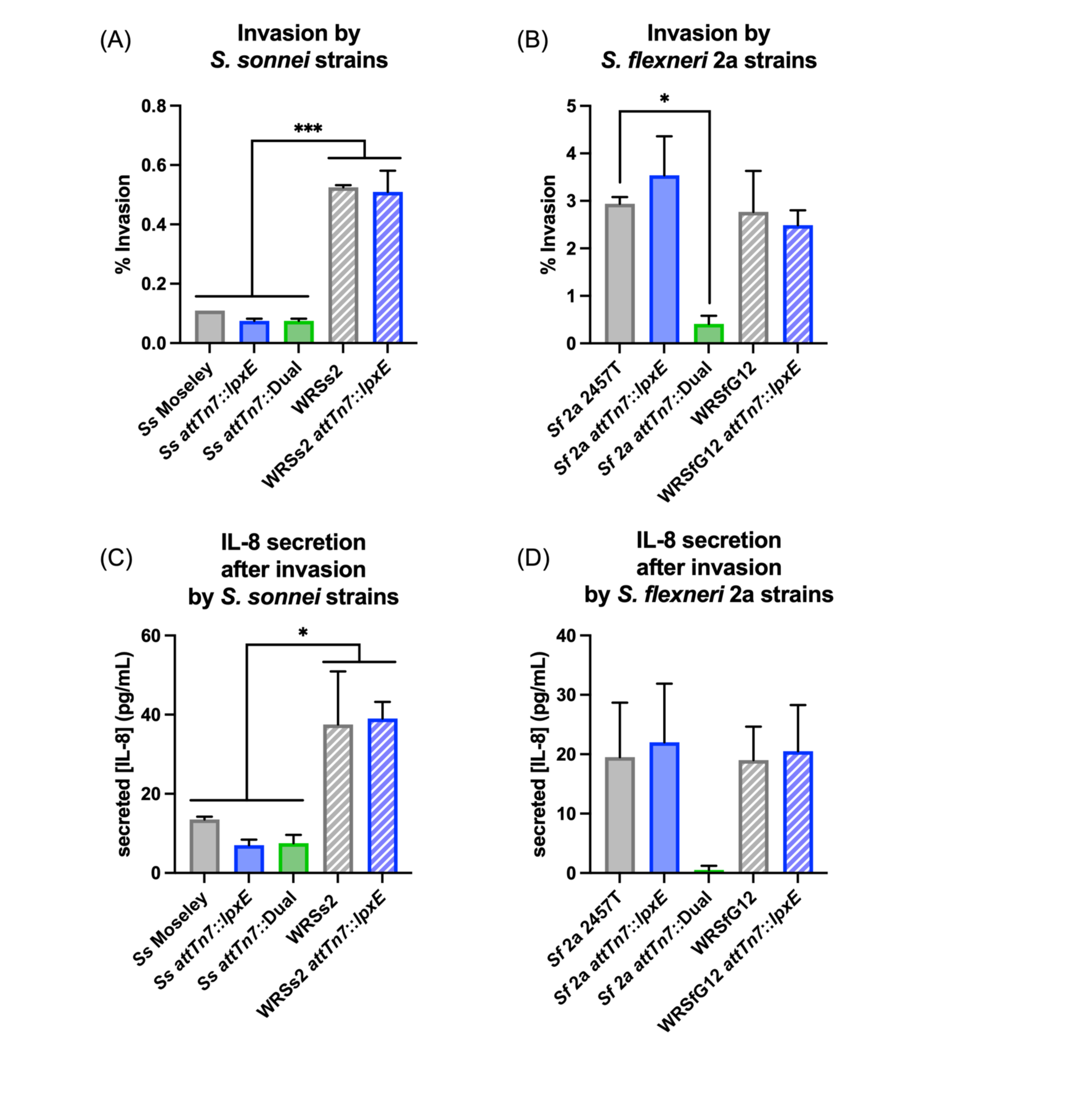
Invasion of HT29 cells by *Shigella* and corresponding IL-8 secretion. (A, B) Invasion of HT29 cells after 4 hours of exposure to an MOI of 10 of *S. sonnei* and *S. flexneri* 2a unmodified and BECC-modified strains. (C, D) IL-8 secretion in the cell supernatant after 4 hours of exposure. Statistical significance determined by ordinary one-way ANOVA. * and *** represent p-values of ≤ 0.05 and ≤ 0.001, respectively.

### Lipid A modifications reduced LPS-induced TLR4 signaling *in vitro*

To assess the TLR4 potency of the BECC-modified LPS, NF-*κ*B reporter cell monolayers were stimulated with increasing amounts of standardized LPS solutions^48,49^. These solutions were normalized, irrespective of modifications, by the quantity of Kdo (2-keto-3-deoxyoctonate) which is a conserved sugar in the inner core region of LPS. Stimulation profiles were fitted with an EC50 value (concentration of LPS that generates half-maximal NF-*κ*B response) where higher EC50 values indicated weaker TLR4 signaling. We found that LpxE- and Dual-modified LPS resulted in diminished NF-*κ*B activation in both murine and human HEK-Blue reporter cells (Figure S5) which corresponded to increased EC50 values compared to unmodified LPS (Table S1). These preliminary results suggested that LpxE- and Dual-modified lipid A variants reduced LPS-mediated TLR4 stimulation to a larger extent than PagL modification. Therefore, chromosomal insertion of P*_ompC_*-*pagL* was not pursued. Purified LPS extracted from chromosomally modified LpxE and Dual BECC-constructs resulted in lower NF-*κ*B activation across several reporter cell lines (Figure 5, S6) and a marked increase in EC50 (Table 1) compared to their parental strains. In some cases, the EC50 was higher in the chromosomally integrated constructs than with the plasmid-based expression (Table 1 vs. Table S1) suggesting increased stability in the chromosomal constructions.

**Figure 5:**
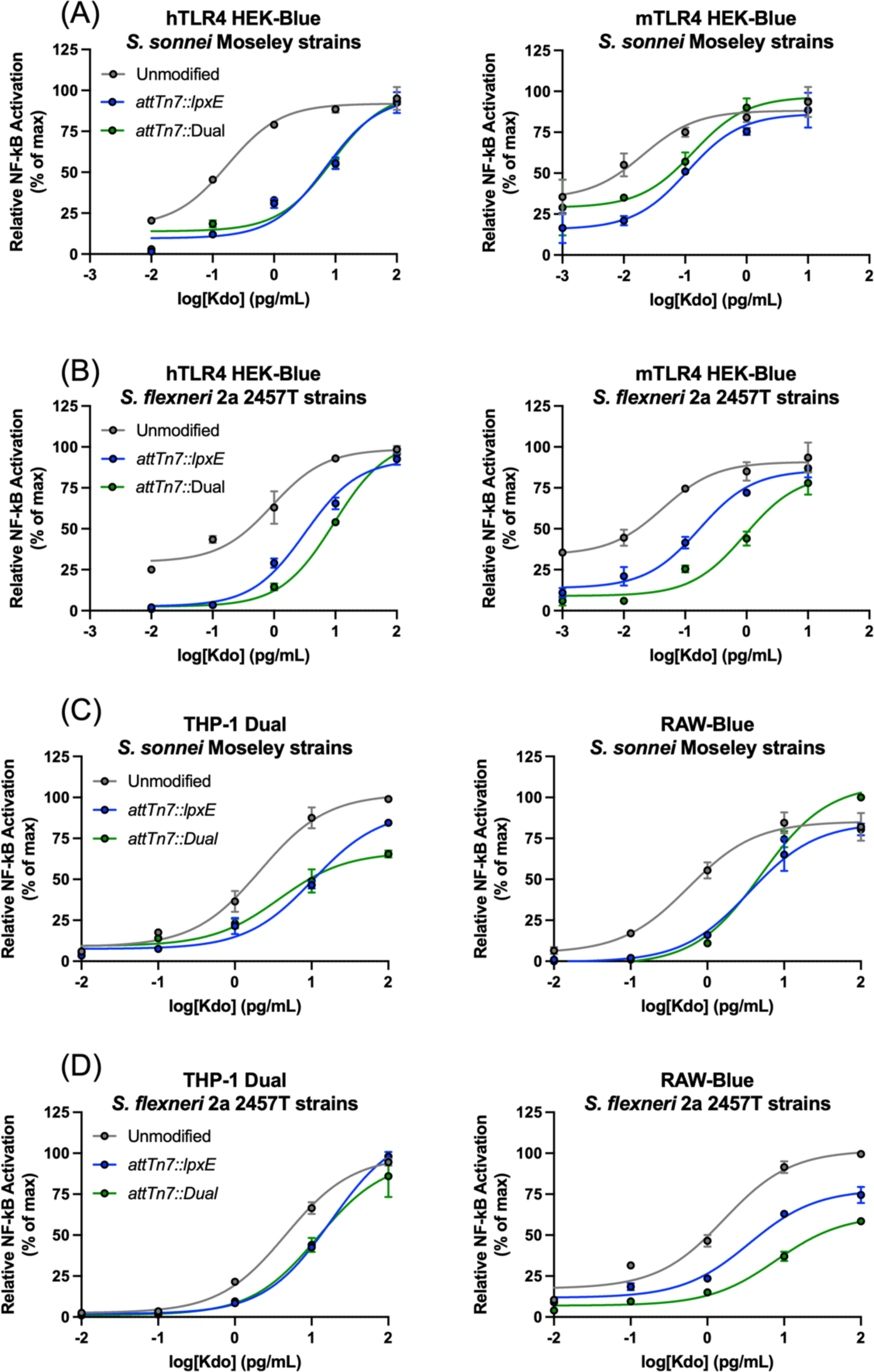
Stimulation of NF-*κ*B reporter cells with LPS solutions from virulent *Shigella* strains. Human(hTLR4)- and mouse(mTLR4)-HEK-Blue, RAW-Blue, and THP-1 Dual cells were stimulated across 10-fold dilutions, in duplicate, of Kdo standardized LPS for 18 hours at 37°C with 5% CO_2_ with purified LPS from wild-type strains of (A, C) *S. sonnei* Moseley and (B, D) *S. flexneri* 2a 2457T.

**Table 1:**
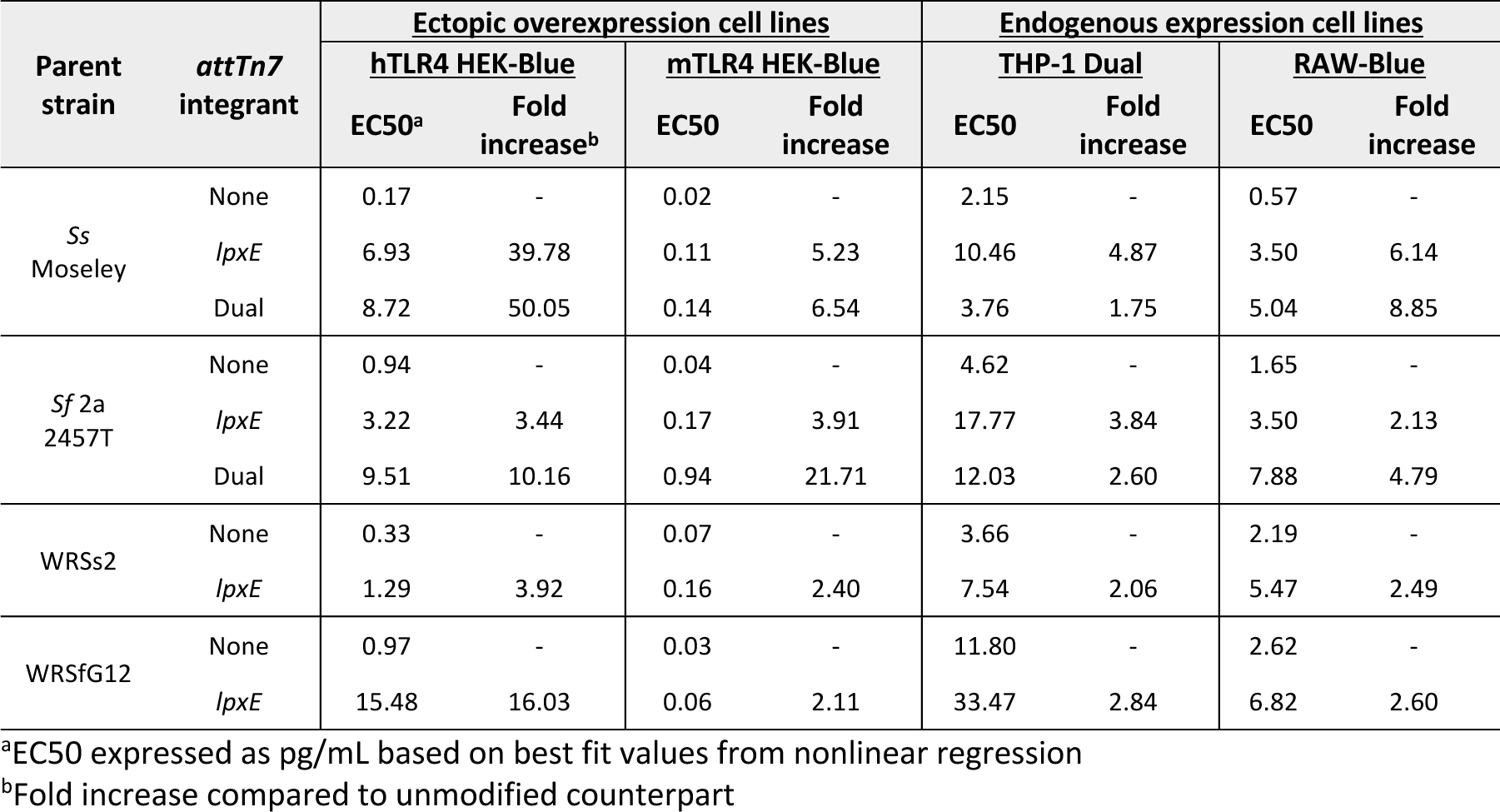
EC50 values for stimulations of NF-*κ*B reporter cells with purified LPS from *attTn7* integrants of the *Shigella* strains used in this study.

| Parent strain | *attTn7* integrant | Ectopic overexpression cell lines |  |  |  | Endogenous expression cell lines |  |  |  |
| --- | --- | --- | --- | --- | --- | --- | --- | --- | --- |
|  |  | hTLR4 HEK-Blue |  | mTLR4 HEK-Blue |  | THP-1 Dual |  | RAW-Blue |  |
|  |  | EC50<sup>a</sup> | Fold increase<sup>b</sup> | EC50 | Fold increase | EC50 | Fold increase | EC50 | Fold increase |
| *Ss* Moseley | None | 0.17 | - | 0.02 | - | 2.15 | - | 0.57 | - |
|  | *lpxE* | 6.93 | 39.78 | 0.11 | 5.23 | 10.46 | 4.87 | 3.50 | 6.14 |
|  | Dual | 8.72 | 50.05 | 0.14 | 6.54 | 3.76 | 1.75 | 5.04 | 8.85 |
| *Sf* 2a 2457T | None | 0.94 | - | 0.04 | - | 4.62 | - | 1.65 | - |
|  | *lpxE* | 3.22 | 3.44 | 0.17 | 3.91 | 17.77 | 3.84 | 3.50 | 2.13 |
|  | Dual | 9.51 | 10.16 | 0.94 | 21.71 | 12.03 | 2.60 | 7.88 | 4.79 |
| WRSs2 | None | 0.33 | - | 0.07 | - | 3.66 | - | 2.19 | - |
|  | *lpxE* | 1.29 | 3.92 | 0.16 | 2.40 | 7.54 | 2.06 | 5.47 | 2.49 |
| WRSfG12 | None | 0.97 | - | 0.03 | - | 11.80 | - | 2.62 | - |
|  | *lpxE* | 15.48 | 16.03 | 0.06 | 2.11 | 33.47 | 2.84 | 6.82 | 2.60 |

<sup>a</sup>EC50 expressed as pg/mL based on best fit values from nonlinear regression
<sup>b</sup>Fold increase compared to unmodified counterpart

### Lipid A modifications diminish LPS-induced cytokine production *ex vivo* from human PBMCs

Preliminary data using LPS extracted from *S. sonnei* Moseley revealed significantly lower TNF-α production upon stimulation of human PBMCs from 6 independent adult donors with BECC-modified LPS when compared to stimulation with unmodified LPS from wild-type Moseley (Figure S7). These results prompted us to stimulate human PBMCs from 4 independent donors with LPS extracted from all strains used in this study and perform multiplex cytokine analysis. In general, all LPS solutions generated similar cytokine induction profiles to each other, irrespective of BECC-modification (Figure 6) but the quantity of cytokine production varied, with BECC-modified LPS having a less stimulating effect than unmodified LPS (Table 2). The most prevalent cytokines, IL-8, IL-6, IFN*γ*, IL-1β, TNF-α, and IL-10 (Table 2 and Figure S8) were present in decreasing abundance, respectively. Additionally, IL-2, IL-4, IL-12p70, and IL-13 were also produced, but in lesser abundance (< 100 pg/mL of supernatant).

**Figure 6:**
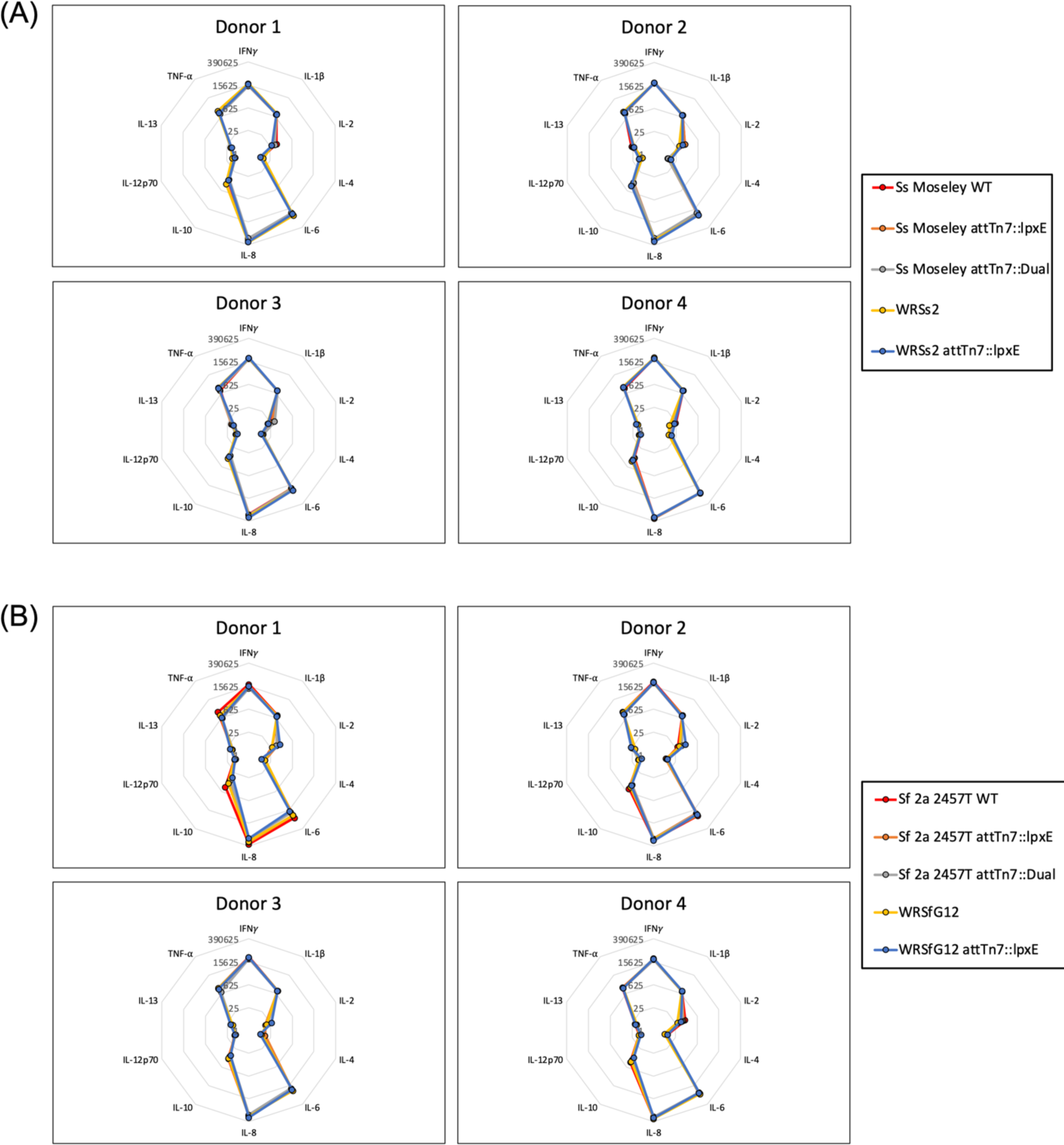
Cytokine induction profiles of stimulated human PBMCs with purified LPS from parental and BECC-modified *S. sonnei* and *S. flexneri* 2a. Human PBMCs from 4 independent donors were stimulated at 1 pg/mL Kdo standardized LPS for 48 hours at 37°C with 5% CO_2_ with purified LPS samples from (A) *S. sonnei* and (B) *S. flexneri* 2a strains used in this study. Cytokine secretion in the supernatant was quantified by MSD multiplex analysis. Cytokine concentrations are expressed as pg/mL via radar plots. Each ring of the plot represents a 25-fold increase in concentration.

**Table 2:** Cytokine levels upon stimulation of human PBMCs with unmodified and BECC-modified LPS.

| Parent strain | LPS from *attTn7* integrant | [Cytokine] (pg/mL) |  |  |  |  |  |
| --- | --- | --- | --- | --- | --- | --- | --- |
| | | TNF-α | IFNγ | IL-1β | IL-6 | IL-8 | IL-10 |
| *Ss* Moseley | None | 440 | 33,340 | 2,563 | 36,382 | 191,316 | 169 |
|  | *lpxE* | 479 | 28,458 | 2,600 | 37,860 | 190,519 | 117 |
|  | Dual | 457 | 26,423 | 2,705 | 37,033 | 169,708 | 126 |
| *Sf* 2a 2457T | None | 1,005 | 40,900 | 4,275 | 46,175 | 220,659 | 224 |
|  | *lpxE* | 584 | 31,130 | 2,770 | 38,933 | 184,402 | 140 |
|  | Dual | 424 | 24,855 | 2,544 | 32,426 | 156,452 | 139 |
| WRSs2 | None | 739 | 34,525 | 3,693 | 45,862 | 215,703 | 186 |
|  | *lpxE* | 612 | 35,875 | 3,305 | 43,339 | 233,600 | 148 |
| WRSfG12 | None | 728 | 33,725 | 3,420 | 44,171 | 198,348 | 209 |
|  | *lpxE* | 475 | 40,323 | 2,349 | 33,837 | 188,210 | 130 |

### Lipid A modifications reduced endotoxicity *in vivo*

To assess if the BECC modifications influenced LPS endotoxicity, mice (n=10) received intraperitoneal injections of Kdo_2_ standardized LPS and were monitored over the course of 72 hours. Clinical scores ranging from 0 – 6 (Table S6) were assigned to each mouse and euthanasia performed if a score above 4 recorded. LPS from unmodified *Shigella* resulted in complete lethality, whereas injections of the same dose of BECC-modified LPS resulted in 100% survival (Figure 7). Mice who received unmodified LPS had clinical scores of 5 by the 24-hour time point, requiring euthanasia. Alternatively, mice who received BECC-modified LPS had clinical scores of 4 or below at the 24-hour time point and recovered to their healthy state (clinical score of 0) by the 72-hour time point. Mice receiving PBS injections recorded clinical scores of 0 throughout the entire study (Figure S9).

**Figure 7:**
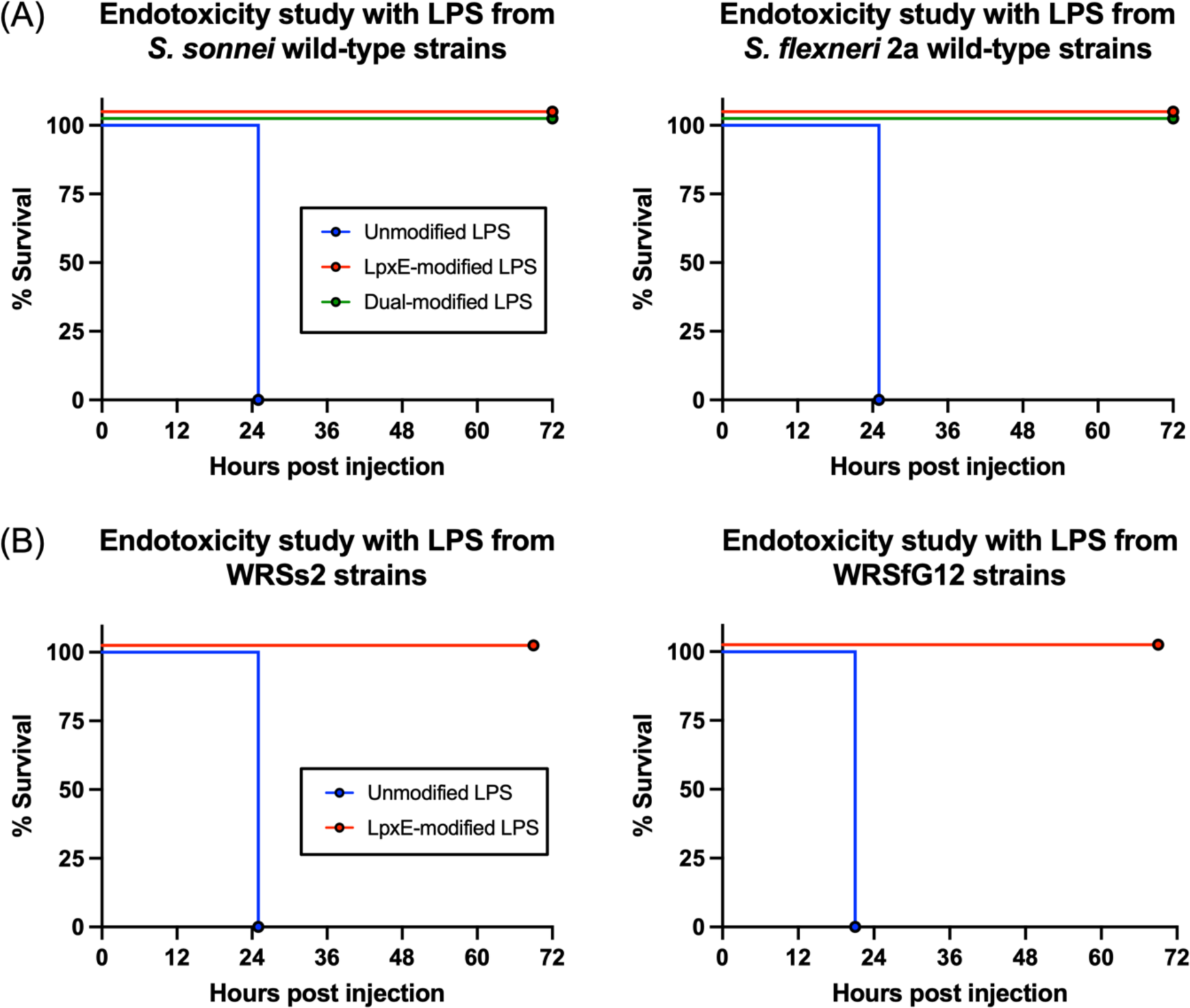
Acute endotoxemia model to assess endotoxicity of LPS from modified *Shigella*. Groups of 10 mice received intraperitoneal injections of a Kdo_2_ normalized dose of LPS, representative of 15 mg/kg. (A) Survival rates for mice receiving purified LPS from unmodified, LpxE-modified, and Dual-modified wild-type strains of *S. sonnei* Moseley and *S. flexneri 2a* 2457T. (B) Survival rates for mice receiving purified LPS from unmodified and LpxE-modified WRSs2 and WRSfG12 strains.

### LpxE-modification did not compromise immunogenicity of the *Shigella* vaccine strains

To evaluate their potential use as vaccine candidates, we measured the immunological response of our BECC-modified *Shigella* vaccine strains in a mouse model. Immunological readouts included serum antibody titers as well as antibody secreting cells (ASC) or cytokine induction profiles in mouse spleen and lung cells to serotype-specific LPS. Initial small scale (< 5 mice/group) vaccine studies employing different routes of vaccination were undertaken to evaluate immunogenicity models. These included oral gastric (o.g.) or intranasal (i.n.) administration of live bacteria (*S. sonnei* WRSs2) as well as i.n. or intramuscular (i.m.) administration of purified *S. sonnei* Moseley LPS (Figure S10A). Vaccinations were delivered two weeks apart using a prime, boost, boost vaccine model. Sera was collected prior to each vaccination and again at day 42 and 56 post-initial vaccination. None of the mice injected intramuscularly with wild-type *S. sonnei* LPS developed an immune response as reflected by the absence of any antibody response against homologous LPS (data not shown). Administration of purified LPS i.n. generated serum IgG antibodies against wild-type *S. sonnei* LPS but no serum IgA antibodies were produced (Figure S10B). Both o.g. and i.n. administration of WRSs2 generated IgG and IgA antibody titers; however, i.n. inoculation was the most favorable route of administration imparting the highest geometric mean serum IgG and mucosal IgA titers (Figure S10B). Increasing the inoculum CFU did not significantly raise antibody production, therefore the lower dose of 10^6^ CFU was utilized for future studies.

Using the data gleaned from the preliminary study above, two large-scale (n=15) independent i.n. vaccination studies were conducted comparing 10^6^ CFU inoculums of the vaccine strains WRSs2 and WRSfG12 to their LpxE-modified variants. To assess markers of the humoral response, total serum IgA and IgG as well as IgG subtypes IgG1 and IgG2a were measured. As shown is Figure 8, vaccination with both WRSs2 *attTn7*::*lpxE* and WRSfG12 *attTn7*::*lpxE* elicited strong serum IgG and IgA immunoglobulin responses against serotype-specific LPS, that mimicked the response from their isogenic parental strain. The combined results of the two vaccination experiments are shown for each mouse in Figure S11. In general, mice exhibited a robust rise in total IgG, IgG1 and IgG2 titers after the first boost. Moderate levels of IgA were also observed. Although there were some significant differences in antibody titers observed between the parental and LpxE-modified vaccine strains at specific time points, only total levels of IgG (WRSs2 day 14) (Figure S11A) and IgG2a and IgA levels (WRSfG12, day 28) (Figure S11B) were statistically significant across both vaccine studies. In addition to serum antibodies, frequencies of total IgG and IgA antibody secreting cells (ASCs) from mouse spleen and lung single-cell suspensions in response to stimulation with serotype-specific LPS were measured as an additional marker of the humoral response to vaccination. Geometric means of total ASC counts for both IgG and IgA were comparable between the parental and LpxE-modified vaccine strains in both vaccine studies (Table S2).

**Figure 8:**
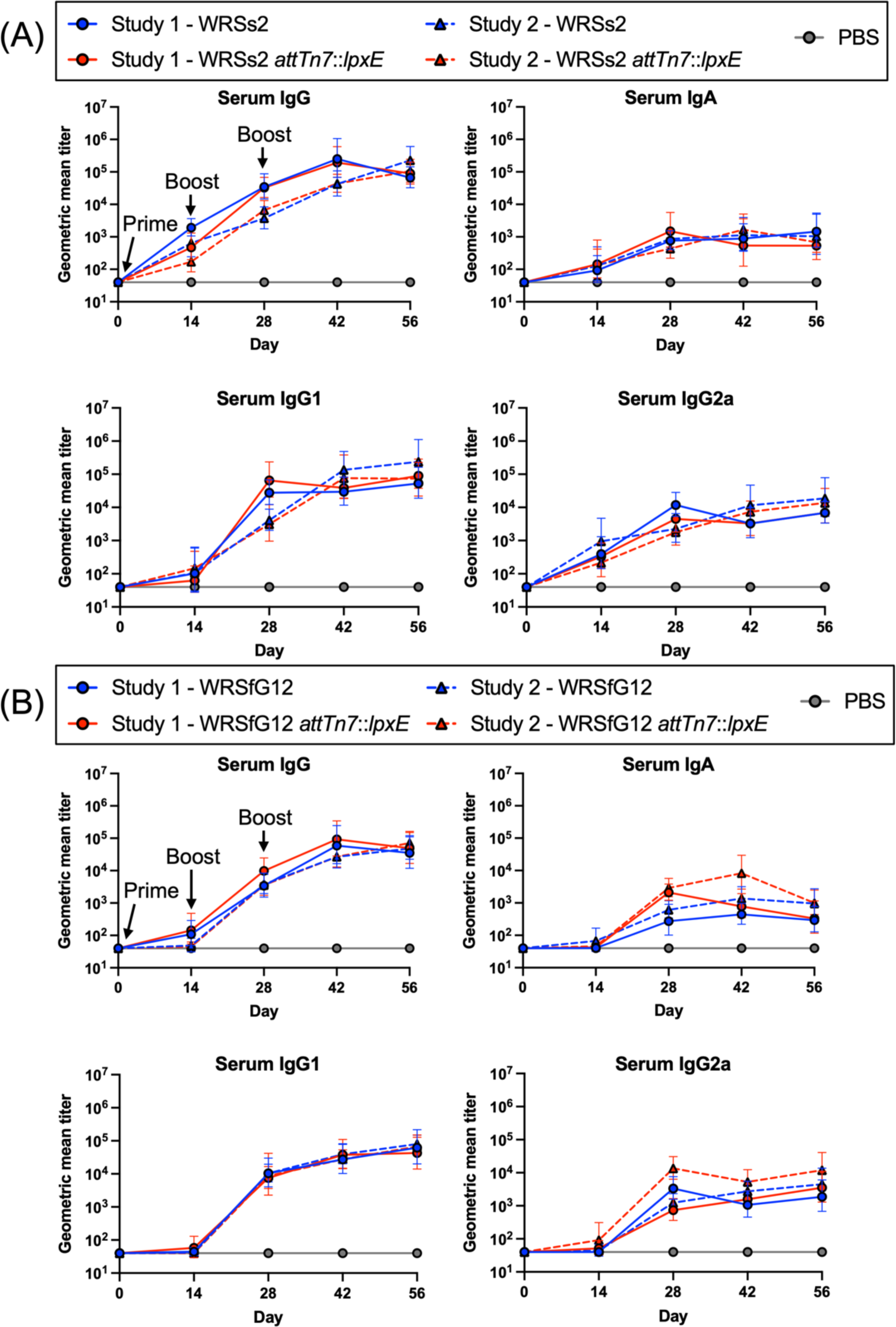
Geometric mean antibody titers from two-independent murine vaccine studies. Serum IgG, IgG1, IgG2a, and IgA antibody titers for each group of mice (n=15) against LPS from wild-type (A) *S. sonnei* Moseley or (B) *S. flexneri* 2a 2457T. Arrows indicate intranasal vaccine administration. Solid lines and circles represent study 1; dashed lines and triangles represent study 2. PBS was used as a control in all studies (grey line and grey circles).

To assess cell-mediated immunity, spleen and lung single-cell suspensions were stimulated with serotype-specific LPS and TNF-α, IL-6, KC/GRO, IL-10, IFN*γ*, IL-1β, IL-2, Il-4, IL-5, and IL-12p70 cytokine profiles were measured via multiplex analysis. Cytokines TNF-α, IL-6, KC/GRO, and IL-10 were produced at high levels (>100 pg/mL) or were increased above basal cytokine levels upon antigen stimulation whereas IFN*γ*, IL-1β, IL-2, Il-4, IL-5, and IL-12p70 were not induced at high levels (< 100 pg/mL) or were at similar concentrations to basal cytokine levels. Regarding the four major cytokines produced at high levels (TNF-α, IL-6, KC/GRO, and IL-10), unvaccinated controls (PBS group) displayed variable results upon exposure to either *S. sonnei* Moseley LPS (Figure S12) or *S. flexneri* 2a 2457T LPS (Figure S13), generating significantly different levels (p ≤ 0.05) of these cytokines in the first vaccine study compared to the second vaccine study in some cases (Figure S12E-H, S13F, H). Vaccinated mice with the parental or LpxE-modified WRSs2 strains displayed no significant difference in production of these cytokines from their isolated spleen and lung cells (Figure S12). Despite, higher cytokine production from LpxE-modified WRSs2 lungs compared to the parental WRSs2 lungs in the first study, this was not repeated in the second study as both the parental and LpxE-modified WRSs2 lungs generated comparable cytokine production (Figure S12E-H). Additionally, vaccinated mice with the parental or LpxE-modified WRSfG12 strains displayed no significant difference in production of these cytokines from their isolated spleen and lung cells(Figure S13). This trend was conserved across both vaccine studies. Taken together, these data demonstrate that the immunogenicity of the vaccine strains was not compromised by modifying the lipid A moiety in the outer membrane.

## Discussion

Despite significant advances in our understanding of *Shigella* pathogenesis and the development of many candidate vaccines^19^, to date, there is no licensed vaccine available. Walter Reed Army Institute of Research has developed live attenuated oral *Shigella* vaccines in both *S. sonnei* and *S. flexneri* 2a which were generally well tolerated but still exhibit unacceptable reactogenicity^31,35^. Here we describe the engineering and characterization of two live attenuated oral *Shigella* vaccines that have been further modified with the promise of reduced reactogenicity and preserved immunogenicity.

Immunity against shigellosis is serotype-specific, suggesting the O-antigen region of LPS is a key antigen for protection^19^. The highly immunostimulatory hexa-acylated bis-phosphorylated lipid A moiety found in *Shigella* LPS induces downstream signaling upon binding TLR4/MD-2, which in turn activates NF-*κ*B and the proinflammatory cytokine cascade^40,43^. Structure activity studies comparing lipid A structure to its ability to signal through TLR4/MD-2 has shown that acylation and phosphorylation drive the magnitude of downstream signaling^39,50–53^. We employed Bacterial Enzymatic Combinatorial Chemistry (BECC), a patented system developed by our laboratory to reduce the toxic effects of LPS. BECC is universally applicable in all Gram-negative bacterial backgrounds and has been previously utilized to modify the lipid A in *Yersinia pestis*, generating well defined variants that have shown success as vaccine adjuvants^54–59^. In this study we used a modified version of *lpxE* isolated from *Francisella* and *pagL*, a gene isolated from *Salmonella*. PagL efficiently removes 3OH C10 acyl chains from the 3-position of lipid A^60^ and LpxE executes dephosphorylation at the 1-position of lipid A^61^. These enzymes were used individually or in combination to either remove a phosphate, an acyl chain or both within the lipid A moiety of the *Shigella* LPS.

When BECC enzymes were expressed from plasmids, multiple lipid A related spectral peaks (Figure S2) indicative of both unmodified and modified species present in the membrane. This incomplete modification of lipid A could be a consequence of plasmid instability even under antibiotic selective pressure. We purified LPS using the standard double hot phenol extraction method^62–64^. Quantification of LPS by dry weight generated variable results in the NF-*κ*B reporter assays, whereas quantitation of Kdo residues contained within the inner LPS core improved consistency, particularly from the plasmid-based expression LPS preparations of BECC-modified *S. sonnei* and *S. flexneri* 2a (data not shown). We therefore used this quantification method throughout our experiments. Using plasmid-expressed BECC constructs in wild-type strains, we first tested the ability of LPS to stimulate NF-*κ*B production *in vitro*. We found that PagL-modification alone had little effect on LPS-induced TLR4 signaling, whereas LpxE- and Dual-modifications led to robust decreases in signaling (Figure S5). This suggested that dephosphorylation was more effective in diminishing LPS-induced TLR4 signaling than deacylation and so we focused our subsequent studies on the LpxE and Dual (LpxE-PagL) modifications. Expression of *lpxE* and Dual cassettes from the chromosome generated spectra with a predominant single base peak (Figure 2), suggesting complete alteration of lipid A inventory on the bacterial membrane.

Two methods, measurement of NF-*κ*B production in cell monolayers and the murine acute endotoxemia model^65^ were used to evaluate reactogenicity of the chromosomally integrated BECC constructs. We used several murine and human NF-*κ*B reporter cell lines and in all experiments, LPS from the LpxE and Dual-modified constructs stimulated less NF-*κ*B than the wild-type LPS and in many cases, there was little difference between the LpxE and Dual-modified constructs (Figure 5). This reduced reactogenicity was also seen in the second-generation vaccine strains WRSs2 and WRSfG12; integration of *lpxE* alone was sufficient to greatly reduce LPS-mediated TLR4 signaling (Figure S6). Previous studies have shown that LPS molecules can signal differentially through different orthologs of TLR4/MD-2^66,67^. These data showed that BECC-modification consistently reduced LPS-mediated TLR4 signaling in both human and mouse orthologs of the TLR4/MD-2 receptor complex. Another method used to evaluate reactogenicity was the murine acute endotoxemia model^65^. In this model, acute endotoxemia is induced by intraperitoneal injection of purified LPS solutions from strains of interest. We found 15 mg/kg which corresponded to injection of 100 µL of 45 µg/mL Kdo_2_ was the optimal lethal dose. Whereas injection of unmodified *Shigella* LPS resulted in complete lethality, injection of the same dose of LpxE- or Dual-modified LPS from our *Shigella* constructs used in this study resulted in 100% survival (Figure 7). These data revealed that the LpxE-modification alone was enough to abrogate the toxic effects of LPS. This may aid in alleviating the reactogenicity associated with ingestion of high doses of *Shigella* vaccine candidates required for protection against shigellosis in humans during clinical trials.

The introduction of extrinsic alterations to a bacterium can sometimes lead to unexpected results. The gene *htrB* encodes for the intrinsic acyl transferase LpxL which is important for a lipid A biosynthesis^41^. Rossi et. al. showed that in *S. sonnei* a deletion in *htrB* generated lipid A lacking an ester linked C12 fatty acid at the 2’-position, leading to reduced TLR4 signaling. The same mutation in *S. flexneri* 2a, however, led to a compensatory addition of an ester linked C16:1 fatty acid in its place, ablating the reduction in signaling compared to unmodified LPS^44^. An advantage of using our BECC approach is that exogenous lipid A modifying enzymes are introduced without genetic manipulation of any endogenous lipid A biosynthetic enzymes. In our study, this approach avoided any compensatory mechanisms that would revert the lipid A structure back to its highly proinflammatory hexacylated bisphosphorylated structure.

Although MALDI-TOF MS analysis confirmed the dephosphorylation and deacylation of lipid A in our constructs, tracking site-specificity was lacking. Native *Shigella* lipid A contains two phosphates and four 3OH C14 acyl groups and MALDI-TOF MS analysis alone did not suggest where the loss of these groups occurred. By using MALDI-TOF MS/MS, precursor ions *m/z* 1797, 1717, and 1490 were selected to undergo further fragmentation. Ester bonds were preferentially cleaved upon fragmentation of precursor ions in the collision cell, likely due to lower stability compared to amide bonds. Additionally, cross-ring cleavage of glucosamine occurred, which enabled elucidation of site-specific information. Analysis of the precursors fragmented into subspecies allowed us to confirm the structure of the modified lipid A, also showing that the lipid A modifying enzymes used are site specific rather than substrate specific. For example, PagL efficiently removes 3OH C10 acyl chains from the 3-position of lipid A in *Pseudomonas*^60^ whereas in *Salmonella* it removes a 3OH C14 acyl chain instead from that same position^68,69^. We observed this as well in our study. Ectopic expression of LpxE and PagL within *Shigella* removed moieties from their expected sites of lipid A; the 1-position for LpxE and 3-position for PagL (Figure 3).

Multiple genetic manipulations can lead to an increase the bacterial metabolic burden leading to deleterious effects in bacterial fitness. Although we were successful with chromosomal insertion of *lpxE* in all strains, we were unable to construct the BECC-Dual (*lpxE-pagL*) modification in the second-generation vaccine strains WRSs2 and WRSfG12. Hence, we tested our constructs to insure these BECC modifications did not impair functional characteristics from their isogenic precursors. As shown in Figure S3, bacterial growth curves for all strains looked similar. The difference in bacterial cell growth between the strains with the greatest variance, *S. sonnei* WRSs2 and *S. sonnei* WRSs2 *attTn7*::*lpxE,* was less than 2-fold. Another key aspect to *Shigella* immunogenicity is the requirement of the large virulence plasmid for *Shigella* host cell invasion and antigen secretion that correlates with protection^70^. Therefore, we chose isolates that were pigmented red on TSA with Congo red dye and were PCR positive for plasmid marker *ipaB,* affirming the presence of the large virulence plasmid. Invasion of gut epithelia has been shown to be necessary for the development of adaptive immunity^9^. We assessed the ability of the BECC-modified *Shigella* to invade mammalian gut epithelia using the well-established gentamicin protection assay^23,25^. None of the LpxE-modified *Shigella* strains were impaired in the gentamicin protection assay. The Dual-modified *S. flexneri* 2a 2457T had significantly lower bacterial invasion compared to parental 2457T (Figure 4, S4) but this was due to the high frequency loss of the invasion plasmid verified by colonies reverting from red to white upon plating on TSA-Congo Red (data not shown).

To observe the innate immune response in a system most likely to represent the response in humans, cytokine profiles of human PBMCs upon LPS stimulation were assessed. Preliminary data from collaborators using LPS from *S. sonnei* Moseley wild-type, *attTn7::lpxE* and *attTn7::Dual* isogenic constructs showed that upon stimulation of PBMCs from 6 independent healthy adults TNF-α secretion was significantly lower for both LpxE- and Dual-modified LPS compared to unmodified LPS (Figure S7). We then examined purified LPS from all the *Shigella* strains used in this study for ten proinflammatory markers using PBMCs from four different healthy adults. The cytokine response patterns from the parental strains were almost identical to their isogenic lipid A modified counterparts (Figure 6). Inspection of the cytokine quantification levels (Table 2) show reduced levels of TNF-α, IFNγ, IL-1β, IL-6, IL-8, and IL-10 from *S. flexneri* 2a 2457T constructs compared to the wild-type 2457T control. Levels for all the above-mentioned cytokines except IFNγ were lower for BECC-modified WRSfG12 LPS stimulation. Except for IFNγ and IL-8, reduced cytokine levels were also observed using LPS from WRSs2 BECC constructs. However, LPS from wild-type *S. sonnei* Moseley was not as consistent and gave lower than expected values for TNF-α, IL-1β and IL-8 (Table 2). Contrasting TNF-α data observed in Figure S7 and Table 2 are most likely due to independent laboratories using different protocols and separate sources of human PBMCs. Taken together, we found that proinflammatory cytokine responses from human immune cells are diminished upon exposure to BECC-modified LPS compared to unmodified LPS.

An important aspect of the host response is the secretion of IL-8 as a potent chemoattractant to recruit polymorphonuclear cells (PMNs) that help to resolve the *Shigella* infection^9^. Cell monolayer supernatants from the gentamycin protection assays were probed for IL-8 secretion to assure that BECC-modified *Shigella* strains still induced IL-8 secretion. Quantification of IL-8 in cell culture supernatants from the invasion assays revealed similar results when comparing parental to BECC-modified *Shigella* (Figure 4C, D). Dual-modified *S. flexneri* 2a 2457T was the only modification that resulted in lower IL-8 secretion (Figure 4D) owing to the loss of the invasion plasmid (see above). Ultimately it was determined that IL-8 secretion was highly correlated to the invasion of the *Shigella* strains (Figure 4 and S4).

*Shigella* is a human-specific pathogen, and therefore does not cause diarrheal related symptoms in small animal models. A preliminary study testing oral gastric, intranasal, and intramuscular vaccination routes determined that intranasal administration was the best vaccination method to reliably generate immune responses in mice (Figure S10). Since a difference in dose response was not seen between 10^6^ and 10^7^ CFU inoculums, the lower dose was chosen for subsequent experiments. To assess humoral immunity, two independent vaccine studies with 15 mice per group were performed comparing serum IgG and IgA titers of unmodified and LpxE-modified WRSs2 and WRSfG12. No difference in titers was observed across mice vaccinated with LpxE-modified or unmodified vaccine strains. Similar results were observed from both vaccine studies comparing geometric mean serum IgG and IgA titers between parental and LpxE-modified vaccine strains including the IgG1 and IgG2a subtypes, which represent Th2 and Th1 cellular immunity, respectively (Figure 8). Consistent with these results, antibody secreting cells (ASC) from mouse spleen and lung exposed to serotype-specific LPS were similar for both parental or LpxE-modified vaccine strains (Table S2) and cytokine secretion from these cells resulted in comparable cytokine profiles for mice vaccinated with the parental or LpxE-modified vaccine strains (Figure S12, S13). Taken together, this data supports the notion that LpxE-modified *Shigella* vaccines would promote a similar adaptive immune response upon vaccination of humans.

The need for a *Shigella* vaccine is becoming increasingly evident. While it has been proposed that endemic *Shigella* can be controlled with public health efforts, the low infectious dose combined with its capacity to acquire extreme drug resistance has bolstered vaccination as a favorable approach to controlling the spread of this pathogen. A recent rise in infection rates by antibiotic resistant strains within developed countries like the United States^12^ suggests that the spread of *Shigella* infections may no longer be localized solely to those living in or traveling to developing countries. Previous live-attenuated *Shigella* vaccines have been tested in clinical trials with immunogenic success, but adverse reactions have been a developmental obstacle. This study modified the immunostimulatory LPS component of the bacterial membrane by altering its lipid A structure and showed reduced endotoxicity without compromising immunogenicity or invasiveness. The BECC-modified vaccine strains of *Shigella* developed in this study have promise to be safer, better tolerated live-attenuated vaccine candidates.

## Acknowledgements

We would like to thank Dr. David Rasko for analyzing the genomic data to confirm the site-specific location of the chromosomal integration of BECC enzymes. We would also like to thank the Department of Defense Medical Research Program, W81XWH1920025, for funding.

## Author contributions

M.E.S., J.M., S. B., M.V., L.C., and R.K.E. designed the study. M.E.S., J.M., S.B., M.V., L.C., and R.K.E. participated in discussion and interpretation of the results. M.E.S characterized the strains and the LPS biological properties *in vitro* and *in vivo*. J.M. designed the constructs and established the molecular biology techniques. S.D. aided in executing the animal work. H.Y. performed the MS/MS analysis. M.E.S. wrote the manuscript. R. K. E. revised the manuscript.

## Declaration of interests

The authors declare no competing interests.

## Methods

### Ethics statement

All animal procedures were approved by the University of Maryland, Baltimore Institutional Animal Care and Use Committee (IACUC #0222002). In all studies, female BALB/cJ mice (Jax Laboratories) were utilized. Mouse husbandry was conducted according to the procedures established at the University of Maryland, Baltimore.

### Bacterial strains and growth conditions

Bacterial strains used in these studies are listed in Table S3. Bacteria were grown at 30°C or 37°C in Lysogenic Broth (Teknova) and Tryptic Soy broth (TSB) or on Tryptic Soy agar (TSA) (Becton Dickinson) supplemented with neomycin (50 µg/ml) (Sigma) or carbenicillin (60 µg/mL) (Sigma) as needed. All *pagL-*expressing strains were supplemented with 1 mM MgCl_2_ to repress the PhoPQ two-component regulatory system. Growth curves were performed in a flat-bottom 96-well uncoated sterile plate (Costar) and recorded using a Cerillo Stratus instrument (Cerillo). Bacteria from a TSA plate were suspended in phosphate buffered saline (PBS) pH 7.4 (Quality Biologicals) to create a 0.5 McFarland standard (1.5 x 10^8^ CFU/mL). The cell density was adjusted 1 x 10^7^ CFU/mL in PBS and 10 µL was used to inoculate 200 μl of TSB/well. Plates were incubated with shaking (180 RPM), at 37°C for 15 hours. Absorbance readings at 600 nm were taken every 15 minutes.

### Molecular genetic techniques

Standard DNA techniques, liquid media, and agar plates were used as described^71^. Restriction endonucleases and T4 DNA ligase were used as recommended by the manufacturer (New England Biolabs). DNA used for cloning purposes was PCR amplified using 10mM dNTP mix (Thermo Scientific) and high-fidelity DNA polymerases Q5 (New England Biolabs) or Pfu ultra II fusion HS (Agilent) according to manufacturer’s instructions. Go-Taq polymerase (Promega) was used for genetic screening. DNA oligonucleotides were obtained from Integrated DNA Technologies and are listed in Table S4. All plasmid constructs (Table S5) were confirmed by double-stranded sequencing (Azenta) and maintained in *E. coli* DH5α or *E. coli* TOP10 (ThermoFisher).

### Generation of plasmid-based BECC-modified *Shigella* strains

LPS modifying enzymes LpxE, PagL, and LpxE-PagL in tandem (termed “Dual”) were first cloned and expressed in plasmid pSEC10^72^ under the osmotically controlled *E. coli ompC* promoter (P*_ompC_*). A codon optimized form of *lpxE* from *Francisella novicida* was synthesized by GenScript and cloned into pUC57 yielding pUC57::*lpxE*. The 720 bp *lpxE* gene was amplified by PCR from pUC57::*lpxE* using Q5 polymerase (New England Biolabs) and primer set *lpxE*-F/*lpxE*-R, trimmed with restriction enzymes BamHI and NheI, and ligated into the BamHI/NheI site of pSEC10 resulting in the construct pSEC10M::P*_ompC_-lpxE.* The *pagL* gene was amplified by PCR from *Salmonella minnesota* (Genbank accession AE006468.2) using Q5 polymerase and primer set *pagL*-F/*pagL*-R. A 570 bp amplicon was trimmed with BamHI/NheI and ligated into the 6630 bp BamHI/NheI digested fragment of pSEC10 yielding pSEC10M::P*_ompC_*-*pagL*. Both *lpxE* and *pagL*, each preceded by a ribosomal binding site, were synthesized in tandem behind the P*_ompC_* and cloned into vector pUC57K by Genscript yielding pUC57K::P*_ompC_*-Dual. The initial subcloning of *pagL* into pSEC10 included six additional bps (GTGTAT) that encoded an alternative start codon present in the *S. minnesota* sequence; this 6 bp sequence was not included in the Dual construct. The P*_ompC_*-Dual gene cassette was then cloned into a modified version of pSEC10 (pSEC10M). Briefly, primer set pSEC10M-F/pSEC10M-R was self-annealed, trimmed with EcoRI/NheI and ligated into a 5775 bp gel purified EcoRI/NheI digested pSEC10 and transformed into *E. coli* TOP10. Resulting pSEC10M had a multiple cloning site [EcoRI-NotI-SwaI-NheI] in place of the *ompC* promoter and *clyA* gene. The 1.9 kb SwaI P*_ompC_*-Dual gene cassette isolated from pUC57K::P*_ompC_*-Dual was ligated into the SwaI digested site of pSEC10M and transformed into *E. coli* TOP10 yielding construct pSEC10M::P*_ompC_*-Dual. Plasmids pSEC10::P*_ompC_*-*lpxE*, pSEC10::P_ompC_-*pagL*, and pSEC10M::P_ompC_-Dual were electroporated into the wild-type strains of *S. sonnei* and *S. flexneri* 2a and selected on TSA with neomycin 50μg/ml (neo50). Successful transformants were used to inoculate a 2 mL overnight culture in TSB (neo50), which received a final concentration of 15% glycerol, followed by storage at −80°C.

### Generation of *attTn7* chromosomally integrated BECC-modified *Shigella* strains

We mobilized P*_ompC_*-*lpx*E and P*_ompC_*-Dual into the *Shigella* chromosome *att*Tn7 site using a site-specific insertion method utilizing the Tn7 recombination machinery on a temperature sensitive plasmid pGRG36^73^. A 1225 bp amplicon of the P*_ompC_-lpxE* gene cassette was generated using template pSEC10::P*_ompC_-lpxE*, Q5 polymerase (New England Biolabs) and primer set P*_ompC_*-F/*lpxE*-R. This was then blunt-ligated into the SmaI digested site of pGRG36, yielding pGRG36::P*_omp_C*-*lpxE*. A 1.9 kb P*_ompC_*-Dual gene cassette flanked by SwaI restriction sites was isolated from pUC57K::P*_ompC_*-Dual and ligated into the SmaI site of pGRG36 yielding pGRG36::P*_ompC_*-Dual. Resulting pGRG36 constructs were transformed into *E. coli* S17-1 and introduced into wild-type and 2^nd^ generation vaccine strains of *S. sonnei* and *S. flexneri* 2a by conjugal mating. Briefly, 2 ml cultures of *E. coli* S17-1 plasmid transformants supplemented with 60 µg/mL carbenicillin (Cb60) and *Shigella* were grown overnight at 30°C and 37°C respectively. Filter matings were performed by mixing 100 µL *Shigella* with 50 µL *E. coli* S17-1 plasmid transformants, concentrated by centrifugation (8k x g for 1 minute), and a 200 μl TSB resuspension was spread onto a 0.45 µM nylon filter (MSI Magna nylon 66) placed on the center of a TSA plate and incubated for 5 hours at 30°C. The nylon filter resuspended in 1 mL TSB and plated out onto TSA containing 0.01% Congo red dye (Sigma-Aldrich), Cb60, 1 mM MgCl_2_ (when needed) and incubated at 30°C. *Shigella* conjugants that grew at 30°C and were Cb60 resistant were screened by PCR for the presence of *lpxE* and the large virulence plasmid using GoTaq (Promega) and primer sets *lpxE*-F/*lpxE*-R (*lpxE*) and *ospD3*-F/*ospD3*-R (*ospD3*) or *lpxE*-F/*lpxE*-R (*lpxE*) and *ipaB*-F/*ipaB*-R (*ipaB*) for virulent and vaccine strains, respectively. Bacteria were plated on TSA containing 0.1% arabinose and incubating at 42°C overnight to promote Tn7 recombination and simultaneous curing of pGRG36. Isolates that were Cb60 sensitive, and Congo Red positive, were assayed by PCR for the presence of *lpxE* using primer set *lpxE*-F/*lpxE*-R and for the large virulence plasmid using ospD3 primer set *ospD3*-F/*ospD3*-R or *ipaB* primer set *ipaB*-F/*ipaB*-R for virulent and vaccine strains, respectively. Confirmed integrants were stored in 15% glycerol at −80°C. Bacterial genomes were sequenced at the Microbial Genome Sequencing Center (SeqCenter) and analyzed using the RAST software^74^ which confirmed the insertion of the gene cassettes into the *att7* site.

### MALDI-TOF MS and MS/MS analysis of lipid A

Functional screening of *Shigella* strains to confirm lipid A modification was performed using the Fast Lipid Analysis Technique (FLAT) coupled to MALDI-TOF mass spectrometric (MS) analysis ^46^. Briefly, a single colony was spotted on a MALDI plate and overlaid with 1 µL of citrate buffer (200 mM citric acid, 100 mM trisodium citrate, pH 3.5). The plate was incubated in a humidified, closed, glass chamber for 30 minutes at 110°C, cooled, washed with endotoxin free water and 1 µL of 10 mg/mL norharmane matrix (Sigma-Aldrich) dissolved in chloroform : methanol (2:1 v/v) was spotted onto the samples on the MALDI plate. MALDI-TOF MS analysis was performed using a Bruker Microflex LRF equipped with a 337 nm nitrogen laser. Spectra were acquired in the negative ion and reflectron mode. Analyses were conducted at < 60% global intensity with 300 laser shots for each spectrum acquisition. Spectra were recorded in triplicate. Agilent ESI tune mix (Agilent) was used for mass calibration. FlexAnalysis software version 3.4 (Bruker) was used to process the mass spectra with smoothed and baseline corrections. Further structural lipid A characterization was conducted by tandem mass spectrometry (MS/MS) analysis using the FLAT^n^ procedure ^47^. The FLAT process above was repeated, except the colony spotted onto an indium tin oxide (ITO) slide instead of a MALDI plate. MS/MS analysis was performed using a Bruker MALDI trapped ion mobility spectrometry Time-of-Flight (timsTOF) mass spectrometer equipped with a dual ESI/MALDI source with a SmartBeam 3D 10 KHz frequency tripled 355 nm Nd:YAG laser. The system was operated in “qTOF” mode (tims deactivated). Ion transfer tuning used the following parameters: Funnel 1 RF: 440.0 Vpp, Funnel 2 RF: 490.0 Vpp, Multipole RF 490.0 Vpp, is CID Energy: 0.0 eV, and Deflection Delta: −60.0 V. The quadrupole used the following values for MS mode: Ion Energy: 4.0 eV and Low Mass 700.00 m/z. Collision cell activation of ions used the following values for MS mode: Collision Energy: 9.0 eV and Collision RF: 3900.0 Vpp. The precursor ion was chosen by inputting targeted *m/z* values including two digits beyond the decimal point. Typical isolation width and collision energy were set to 4 – 6 *m/z* and 100 – 110 eV, respectively. Focus Pre-TOF used the following values: Transfer time 110.0 µs and Pre pulse storage 9.0 µs. Agilent ESI Tune Mix (Agilent) was used to perform calibration. MALDI parameters in qTOF mode were optimized to maximize intensity by tuning ion optics, laser intensity, and laser focus. All spectra were collected at a laser diameter of 104 µm with beam scan on using 800 laser shots per spot using either 70% or 80% laser power. MS/MS data were collected in negative ion mode. In all cases, a matrix of 10 mg/mL Norharmane dissolved in chloroform : methanol (2:1 v/v) was used. mMass software version 5.5.0 ^75^ was used to process the mass spectra with smoothed and baseline corrections. Identification of all fragment ions were determined based on ChemDraw Ultra version 10.0.

### Lipopolysaccharide (LPS) extraction and purification

LPS was isolated using the double hot phenol method. Briefly, two liters of bacterial culture were harvested by centrifugation and resuspended in 90% phenol : endotoxin free water (1:1 v/v) and incubated at 65°C for 1 hour. After centrifugation, the aqueous phase was isolated from the two-phase solution (repeated three times total and pooled) and dialyzed for 36 hours against deionized water using pre-treated 1 kD MWCO RC tubing (Spectrumlabs.com), followed by flash freezing and lyophilization. The lyophilized product was resuspended in 20 mM Tris-HCl pH 8.4 supplemented with 2 mM MgCl_2_ and digested using 500 units of Benzonase and 100 µg/mL Dnase I for 2 hours at 37°C. The pH was subsequently adjusted to 7.4 using 1 N HCl and the solution further digested with 100 µg/mL Proteinase K for 2 hours at 37°C. Water-saturated phenol was added, vortexed, centrifuged (8,000 x g) and the upper aqueous phase collected, dialyzed and lyophilized. Further isolation of LPS was performed by serial washes in chloroform : methanol (2:1 v/v) as described ^62^. The LPS was separated from contaminating lipoproteins as described ^63^ by resuspension in 0.2% TEA and 0.5% DOC, followed by the addition of 37°C water-saturated phenol and the upper aqueous phase collected. Finally, the LPS product was precipitated by the addition of cold 100% ethanol and 30 mM sodium acetate followed by incubation for 18 hours at −20°C. The LPS precipitate was harvested by centrifugation (5,000 x g, 20 minutes), washed in cold 100% ethanol, resuspended in endotoxin free water (Quality Biological) and lyophilized.

### Bligh Dyer total lipid extraction

Harvested bacterial pellets from log-phase cultures were resuspended in 1 mL of methanol : chloroform : water (2:1:0.8 v/v/v) and incubated shaking (180 RPM) for 3 hours at room temperature. Next, 500 µL of chloroform : water (1:1 v/v) was added, vortexed, and incubated with shaking for 30 minutes at room temperature. The solution was centrifuged (5,000 x g for 5 mins) and the lower chloroform layer containing LPS was isolated. To the cell pellet, 500 µL of chloroform was added and incubated with shaking for 30 minutes at room temperature. Following centrifugation (5,000 x g for 5 mins), isolated lower chloroform layers were pooled together. Samples were dried under a gentle N_2_ stream of gas, resuspended in 50 µL chloroform, and 1 µL spotted for MALDI-TOF MS analysis using Norharmane matrix.

### Kdo assay for LPS quantification

2-keto-3-deoxyoctonate (Kdo) standards ranging from 12 – 48 µg/mL in endotoxin free water and 1 mg/mL LPS solution solutions were hydrolyzed in 0.018 N sulfuric acid (H_2_SO_4_) at 100°C for 20 minutes, followed by the addition of 25 µL/sample of 9.1 mg/mL periodic acid (H_5_IO_6_) in 0.125 N H_2_SO_4_ and incubated in the dark for 20 minutes. Addition of 50 µL/sample of 2.6% sodium arsenite (NaAsO_2_) in 0.5 N HCl was followed by addition of 250 µL/sample of 0.3% thiobarbituric acid (TBA). Samples were heated at 100°C for 10 minutes, quickly followed by addition of 125 µL/sample of dimethyl sulfoxide (DMSO), and the absorbance measured at 550 nm. The absolute quantification is based on the interpolation of the standard curve provided the Kdo_2_ quantity. Half of the Kdo_2_ quantity, representing Kdo_1_ (referred to as simply “Kdo” in this study), was utilized for normalization.

### Murine acute endotoxemia

LPS solutions of 45 µg/mL Kdo_2_ were prepared in sterile, endotoxin free, PBS (Quality Biology). LPS solutions were transferred to arbitrarily labeled tubes by a third-party observer to ensure blinding to group identifications and avoiding bias in clinical score designations. One hundred µL/mouse of LPS solution was delivered using a slip tip 1 mL syringe attached to a 27-gauge ½ inch needle (Becton Dickinson) via the intraperitoneal (i.p.) route. Mice were monitored for 72 hours post-injection, receiving a clinical score/mouse based on appearance and mobility as described in Table S6. A clinical score of 5 required euthanization as a consequence of no movement, noticeable stress, and an inability to return upright if placed on their side.

### Cell culture media and conditions

RPMI-1640 (Gibco) complemented with 25 mM HEPES, 2 mM glutamine, 10% FBS, 1% Pen-Strep, referred to as cRPMI (complete RPMI), was filter sterilized through a 0.22 µM filter flask and used for primary lung and spleen cells isolated from mice dissections and the THP-1 NF-*κ*B-SEAP reporter cell line (THP-1 Dual, Invitrogen). DMEM (Corning) complimented with 10% FBS, 3.7 g/L sodium bicarbonate, 2 mM glutamine, and 100 Units/mL penicillin-streptomycin, referred to as complete DMEM (cDMEM), was filter sterilized through a 0.22 µM filter flask and used for HT29 cells (courtesy of Dr. Eileen Barry, UMB), murine TLR4 (mTLR4) and human TLR4 (hTLR4) HEK-Blue cells (Invitrogen), and RAW-Blue cells (Invitrogen). All cells were maintained at 37°C with 5% CO_2_.

### NF-*κ*B reporter cell line stimulations

Sterile, cell culture-treated, 96-well flat bottom plates (Costar) were seeded with HEK-Blue, RAW-Blue, or THP-1 Dual reporter cells at 1 x 10^5^ cells/well. THP-1 cells received 100 ng/mL of vitamin D_3_ (Sigma) prior to seeding in wells to enable cell differentiation. Cells were incubated for 18 hours at 37°C with 5% CO_2_ except the THP-1 cells which were incubated for 72 hours to enable differentiation into monocyte-derived macrophages. Five 10-fold dilutions of Kdo standardized LPS ranging from 10^2^ pg/mL to 10^-2^ pg/mL was used to stimulate NF-*κ*B production in cells. Cells were incubated at 37°C with 5% CO_2_ for 18 hours. Detection of NF*κ*B was quantified using Quanti-Blue (QB) reagent prepared according to manufacturer’s protocols (Invitrogen). Briefly, 1 mL of QB buffer was mixed with 1 mL of QB reagent top stock, dissolved into 100 mL total of sterile endotoxin free water. After incubation, 20 µL of cell supernatant was mixed with 180 µL of QB reagent in 96-well plates and incubated until the highest absorbance reading at 630 nm reached 1.5. The percent of relative NF-*κ*B activation was normalized to the maximum OD 630 nm measured. Points were plotted as the mean ± standard deviation of the relative NF-*κ*B activation at each concentration using GraphPad Prism version 9 and fitted using a nonlinear regression of the log(agonist) versus response (three parameters) via the following equation.

### Stimulation of primary Peripheral blood monocytes (PBMCs)

Peripheral blood monocytes (PBMCs) obtained from 4 independent donors (AllCells), were prepared by two washes of warm cRPMI and seeded at 5 x 10^5^ cells/well in 96-well, sterile, uncoated round bottom plates (Costar) in the presence of 1 pg/mL Kdo standardized LPS solutions dissolved in cRPMI. Plates were incubated at 37°C with 5% CO_2_ for 48 hours. Supernatants were recovered after centrifugation (400 x g, 5 minutes) and stored at −20°C until analyzed by MSD multiplex.

### Invasion assays

HT29 cells were seeded at a density of 5 x 10^5^ cells/well in 24-well flat bottom sterile plates (Corning) and incubated at 37°C for 18 hours with 5% CO_2_. *Shigella* cultures from overnight growth on TSA containing 0.01% Congo Red were used to generate a resuspension in sterile PBS pH 7.4. A 5-fold dilution generating an OD_600_ of roughly 0.5 (actual OD_600_ used for calculation) was then used to back calculate estimated CFU/mL using the following equation:

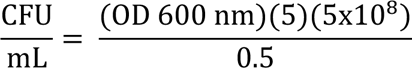

Dilutions of 5 x 10^6^ or 5 x 10^7^ CFU/mL were used for a multiplicity of infection (MOI) of 10 or 100 respectively. Enumeration of the inoculation was confirmed in duplicate by plate counts on TSA. Bacteria were added to media-free, PBS-washed cell monolayers and the plates were centrifuged (3,000 x g, 5 minutes). After incubation for 90 minutes at 37°C with 5% CO_2_ cell monolayers were twice washed with sterile PBS followed by the addition of cDMEM containing 50μg/ml gentamycin (Sigma-Aldrich) and incubated for 2.5 hours. Supernatants were isolated and processed for human IL-8 secretion by cytokine ELISA. The cell monolayer was twice washed with sterile PBS and lysed with 1% Triton X-100 (Sigma-Aldrich) in sterile PBS for 10 minutes at room temperature. Serial dilutions were plated in duplicate on TSA and incubated overnight at 37°C. The percentage of invasion was determined as the CFU/mL recovered normalized to the CFU/mL inoculated.

### Cytokine ELISA

Cytokine analysis of host cell culture supernatants was performed using DuoSet ELISA kits (R&D Systems) according to the manufacturer’s protocol. Briefly, plates were coated overnight at 4°C by adding 100 µL/well of 2 µg/mL capture antibody in ELISA coating buffer, washed three times PBS + 0.02% Tween-20 (PBST) and blocked with 300 µL/well 1% BSA in PBS for 1 hour at room temperature followed by three washes with PBST. Cell culture supernatants were diluted to reach a signal within the dynamic range. Bound cytokines were labeled with by adding biotin-conjugated antibodies in block buffer (100 µL/well of 2 pg/mL) and incubated at room temperature for 2 hours. Plates were washed with PBST, incubated for 20 minutes with secondary antibody streptavidin-HRP followed by addition of a color substrate. The plates were read at both 450 nm and 562 nm and the difference taken as the final reading. The amount of cytokine is reported as picograms per mL of cell culture supernatant.

### Murine vaccination with live-attenuated *Shigella*

*Shigella* vaccine strains WRSs2 and WRSfG12 were grown at 37°C overnight on TSA containing 0.1% Congo Red. <u>Intranasal (i.n.) vaccination</u>: Vaccine inoculums of 3.33 x 10^8^ CFU/mL and 3.33 x 10^7^ CFU/mL of *Shigella* were prepared in sterile PBS using the method and equation above (see invasion assays) and kept at room temperature. Mice were anesthetized using a Matrx VIP 3000 vaporizer (Midmark Animal Health) with isoflurane (Fluriso, VetOne) : oxygen (Airgas, OX USPEAWBDS) mixture (1:1 mixing) for 1-2 minutes and 15 µL of the vaccine inoculum was delivered to each nare (30 µL in total) using a pipette. <u>Oral gastric (o.g.) vaccination:</u> Vaccine inoculums of 1 x 10^8^ CFU/mL and 1 x 10^7^ CFU/mL of *Shigella* were prepared in sterile PBS and kept at room temperature. Inoculums of 100 µL were delivered by intra-gastric gavage using a 2-inch-long plastic feeding needle (VWR) connected to a 1 mL syringe (Becton Dickinson). For both vaccination methods, mice were monitored for adverse reactions post immunization. Enumeration of the vaccine inoculums were determined by plate counts on TSA.

### Murine vaccination with purified LPS

Purified LPS from wild-type *S. sonnei* (Ss) Moseley was obtained as described above (see LPS extraction and purification). <u>Intranasal (i.n.) vaccination:</u> Vaccines containing purified Ss LPS at 1 mg/mL and 0.66 mg/mL dissolved in sterile PBS were delivered i.n. as described above (see murine vaccination with live-attenuated *Shigella*). <u>Intramuscular (i.m.) vaccination:</u> Vaccines containing purified Ss LPS at 0.6 mg/mL and 0.4 mg/mL dissolved in sterile PBS were prepared and stored at room temperature. Mice were immobilized using a restrainer and 50 µL of the vaccine was injected using a 1 mL syringe (Becton Dickinson) into the caudal muscle after disinfecting the area with 70% ethanol. For both vaccination methods, mice were monitored for adverse reactions post immunization.

### Sera collection

Mice were bled via the lateral saphenous vein using petroleum jelly and 27-gauge needles. Blood was collected in microvette 200 Z-gel tubes (Sarstedt) and the sera isolated by centrifugation (10,000 x g, 3 minutes) and stored in sealed uncoated 96-well flat bottom plates (ThermoFischer) at −20°C.

### Enzyme-linked immunosorbent assay (ELISA)

Coating antigens used in ELISAs included purified LPS from wild-type *S. sonnei* (Ss) Moseley or *S. flexneri* 2a (Sf 2a) 2457T. Nunc MaxiSorp plates (ThermoFischer) were coated with 5 µg/mL serotype-specific LPS (Ss LPS for WRSs2 vaccinated mice and Sf 2a LPS for WRSfG12 vaccinated mice) in 100 mM carbonate coating buffer pH 9.6 (sodium bicarbonate/carbonate) and incubated for 3 hours at 37°C. Plates were washed with PBS containing 0.05% Tween-20 (Sigma) (PBST) and blocked with 10% non-fat dry milk powder (Quality Biological) in PBST overnight at 4°C. Sera was diluted in 5-fold increments starting with a 1:50 dilution in PBST and added to the LPS-coated plates and incubated at 37°C for 2 hours. Plates were washed with PBST. Incubation for 1 hour with secondary HRP-conjugated antibodies, goat anti-mouse IgG, IgG1, IgG2a (Southern Biotech) or goat anti-mouse IgA (Invitrogen) was followed by a 15 minute room temperature incubation with 3,3ʹ,5,5ʹ-tetramethylbenzidine (TMB) substrate (BD biosciences) prepared according to manufacturer’s protocol. KPL TMB stop solution (Sera Care) containing 1% HCl was added to each well and the absorbance read at 450 nm. The endpoint titer was determined as the absorbance reading that was equal to the reciprocal dilution required for the signal to match the average blank (PBST alone, no sera). Samples were run in duplicate. Sera from days 28, 42, and 56 of the vaccine study required dilutions of 1:1250 for IgG and IgG1 titers and 1:250 for IgG2a to reach specific endpoints.

### Generation of single cell suspensions from spleen and lung harvests

Mice were sacrificed on day 56 after initial immunization. Spleens and lungs were immediately harvested and placed into sterile RPMI 1640 containing 0.3 mg/mL L-Glutamine (Gibco) and placed on ice. Single cell suspensions were generated by procecessing tissues in gentleMACS C-tubes (Miltenyi Biotec) in a gentleMACS Dissociater (Miltenyi Biotec) according to manufacturer’s protocol. Spleen tissue was bathed in sterile RPMI and lung tissue was bathed in a solution of 86% RPMI 1640: 12% collagenase/hyaluronidase (StemCell): 2% DNAseI (10mg/ml) (v/v/v). Lung tissue had a 20-minute shaking (100rpm) incubation period at 37°C in between dissociation episodes. Single cell suspensions were passed through 70 µM MACS smart strainers (Miltenyi Biotec), media removed by centrifugation (800 RPM, 5 minutes) and 1 mL RBC lysis buffer (Sigma-Aldrich) was added to eliminate any red blood cells. Lysis was halted by the addition of 7 mL RPMI 1640 supplemented with 10% FBS (complete RPMI, cRPMI). Cell suspensions were passed through 70 µM MACS smart strainers, centrifuged (800 RPM, 5 minutes) and the cell pellet was resuspended in 500 µL cRPMI. Cells were diluted 50-fold into RPMI 1640 and counted using a DeNovix CellDrop.

### Antigen stimulation of primary murine cell suspensions

Serotype-specific Kdo standardized LPS antigen (100 pg/mL) and 1 x 10^6^ spleen cells or 1 x 10^5^ lung cells were mixed in uncoated round-bottom sterile 96-well plates (Costar). Plates were incubated at 37°C with 5% CO_2_ for 48 hours. Supernatants were recovered after centrifugation (400 x g, 5 minutes) and stored at −20°C until analyzed by MSD multiplex (Meso Scale Discovery).

### ELISpot analysis of primary spleen and lung cell suspensions

ELISpot plates (Cellular Technology Limited) were performed according to the manufacturer’s protocol. Briefly, the membrane was activated by adding 70% EtOH immediately followed by a PBS wash and blotted to remove excess liquid. 80 µL/well of manufacturer’s capture antibody solution (anti-mouse Ig*κ* and anti-mouse Igƛ) solution was added. The coating solution was decanted, plates washed with PBS, and incubated for 1 hour at room temperature with 150 µL/well cRPMI. Media was replaced with 100 µL/well cRPMI containing a working concentration of 100 pg/mL Kdo standardized LPS followed by 100 µL/well of 1 x 10^5^ cells/well along with two 5-fold dilutions for both spleen and lung. Plates were incubated static for 18 – 24 hours at 37°C with 5% CO_2_. Plates were processed according to manufacturer’s protocols and spots were scanned and counted using an ImmunoSpot Analyzer and ImmunoSpot Software (Cellular Technology Limited). Individual red and blue spots were recorded by hand.

### Multiplex cytokine analysis

MSD (Meso Scale Development) V-PLEX mouse proinflammatory panel 1 (10-plex) was used for the analysis of the mouse orthologs of IFN*γ*, IL-1β, IL-2, IL-4, IL-5, IL-6, KC/GRO, IL-10, IL-12p70, and TNFα from 25 µL of undiluted supernatant from the antigen stimulation of primary murine spleen and lung cell suspensions. MSD V-PLEX human proinflammatory panel 1 (10-plex) was used for the analysis of the human orthologs of IFN*γ*, IL-1β, IL-2, IL-4, IL-6, IL-8, IL-10, IL-12p70, IL-13 and TNFα from 25 µL of 6-fold dilutions of the supernatant from the stimulation of human PBMCs. Samples and calibrators were incubated at room temperature, shaking (500 RPM), for 2 hours. Plates were washed with PBST and MSD detection antibody solution, prepared according to manufacturer’s protocol was added and incubated shaking (500 RPM), at room temperature, for 2 hours. Plates were washed with 150 µL/well PBST followed by the addition of 150 µL/well of read buffer T. Plates were immediately read on an MSD SQ 120/120MM instrument. Cytokine concentration was determined by interpolation from a standard curve generated using the provided calibrators.

## Notes

### Competing Interest Statement

The authors have declared no competing interest.

