## Supplemental information for "Targeted lipid A modification of *Shigella* vaccine strains reduced endotoxicity without compromising immunogenicity or invasiveness"

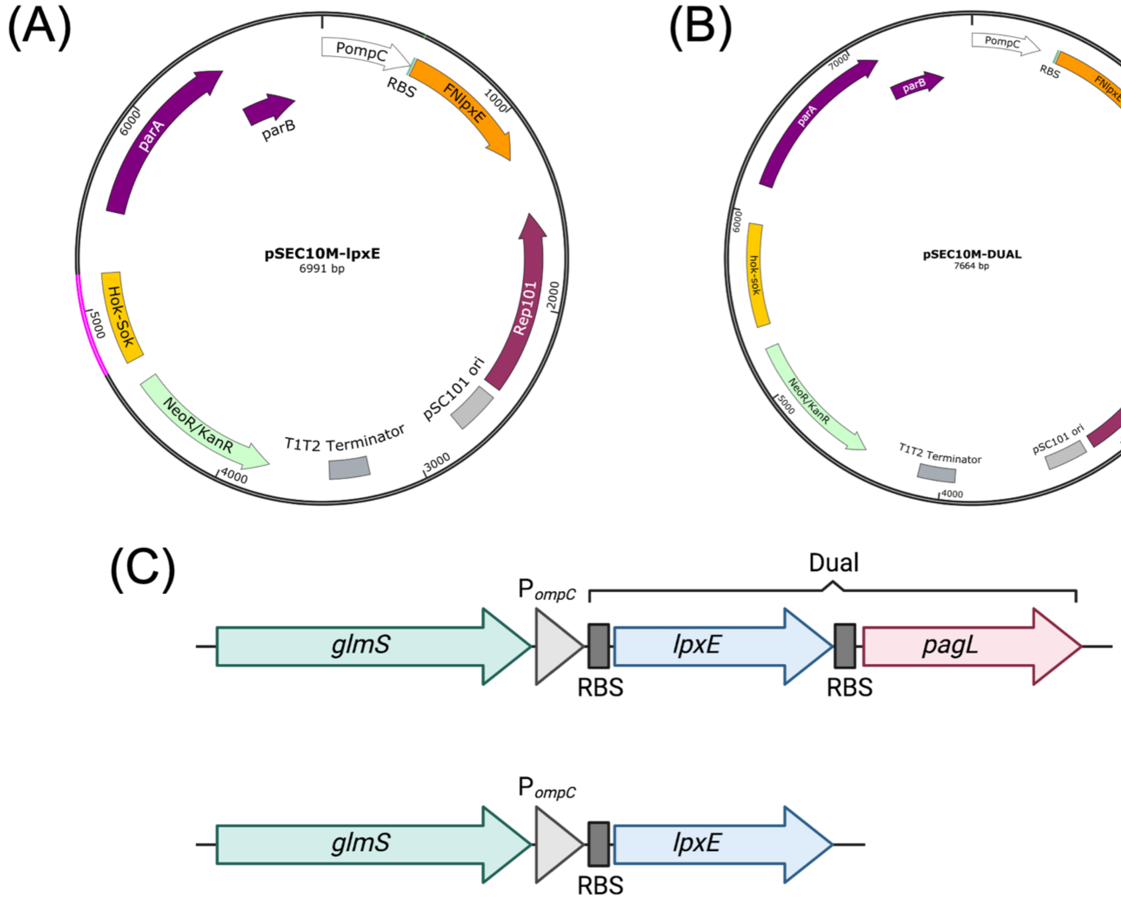

**Figure S1:** Plasmid and chromosomal constructs

For plasmid-based expression, BECC constructs expressed (A) *lpxE* alone or (B) both *lpxE* and *pagL* (termed "Dual") from the osmotically controlled *ompC* promoter ($P_{ompC}$) within the pSEC10M plasmid. For chromosomal-based expression, BECC constructs were expressed from $P_{ompC}$ in the *attTn7* site at the 3' end of the *glmS* gene.

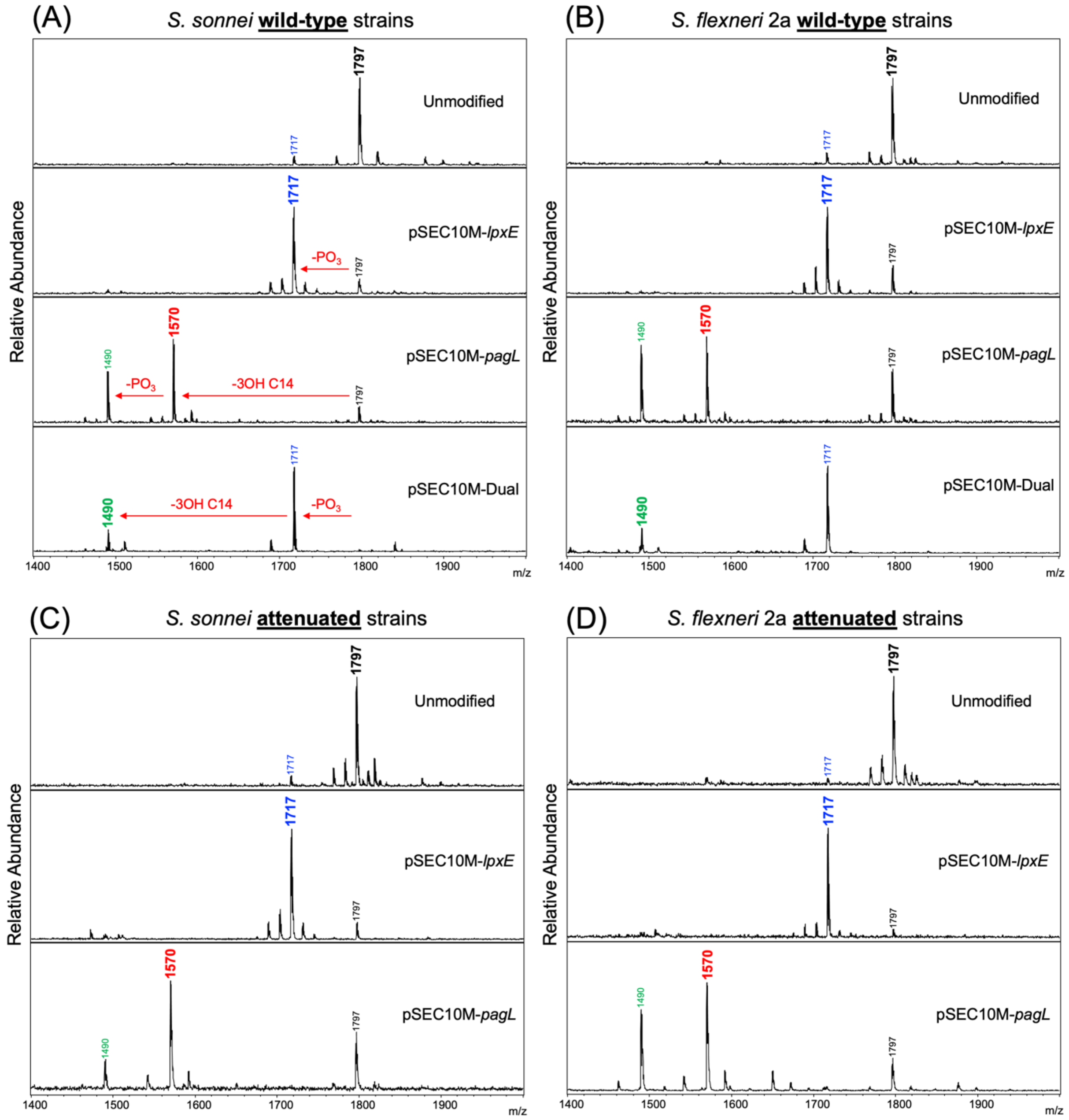

**Figure S2:** MALDI-TOF MS spectra of *Shigella* strains expressing BECC enzymes from plasmids

Representative spectra from wild-type strains of (A) *S. sonnei* Moseley and (B) *S. flexneri* 2a 2457T as well as from attenuated strains (C) WRSs1 and (D) SC602. Spectral peaks represent $[M-H]^-$ ions. Colored peaks correspond to the expected structures detailed in Figure 1. Arrows depicted in (A) indicate the loss of a phosphate ($PO_3$) or acyl chain (3OH C14) from the native lipid A structure.

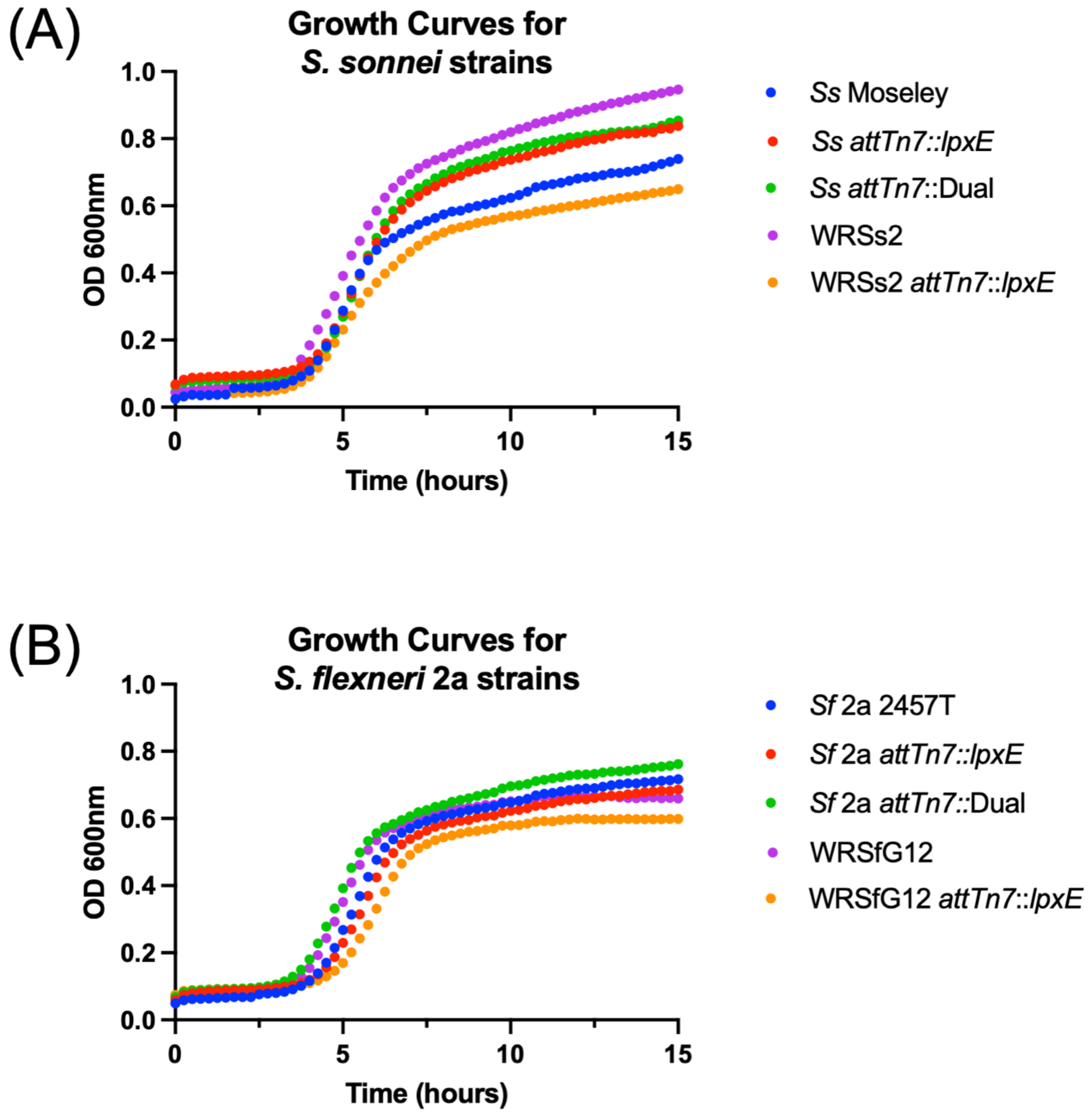

**Figure S3**: Growth curves for *Shigella* strains used in this study

Growth over time, monitored using a Cerillo growth curve instrument, for $10^5$ CFU/mL inoculums in 96-well plates for (A) *S. sonnei* Moseley and (B) *S. flexneri* 2a 2457T strains and their BECC variants. Plotted values are the mean of triplicate readings. Error bars are omitted for clarity.

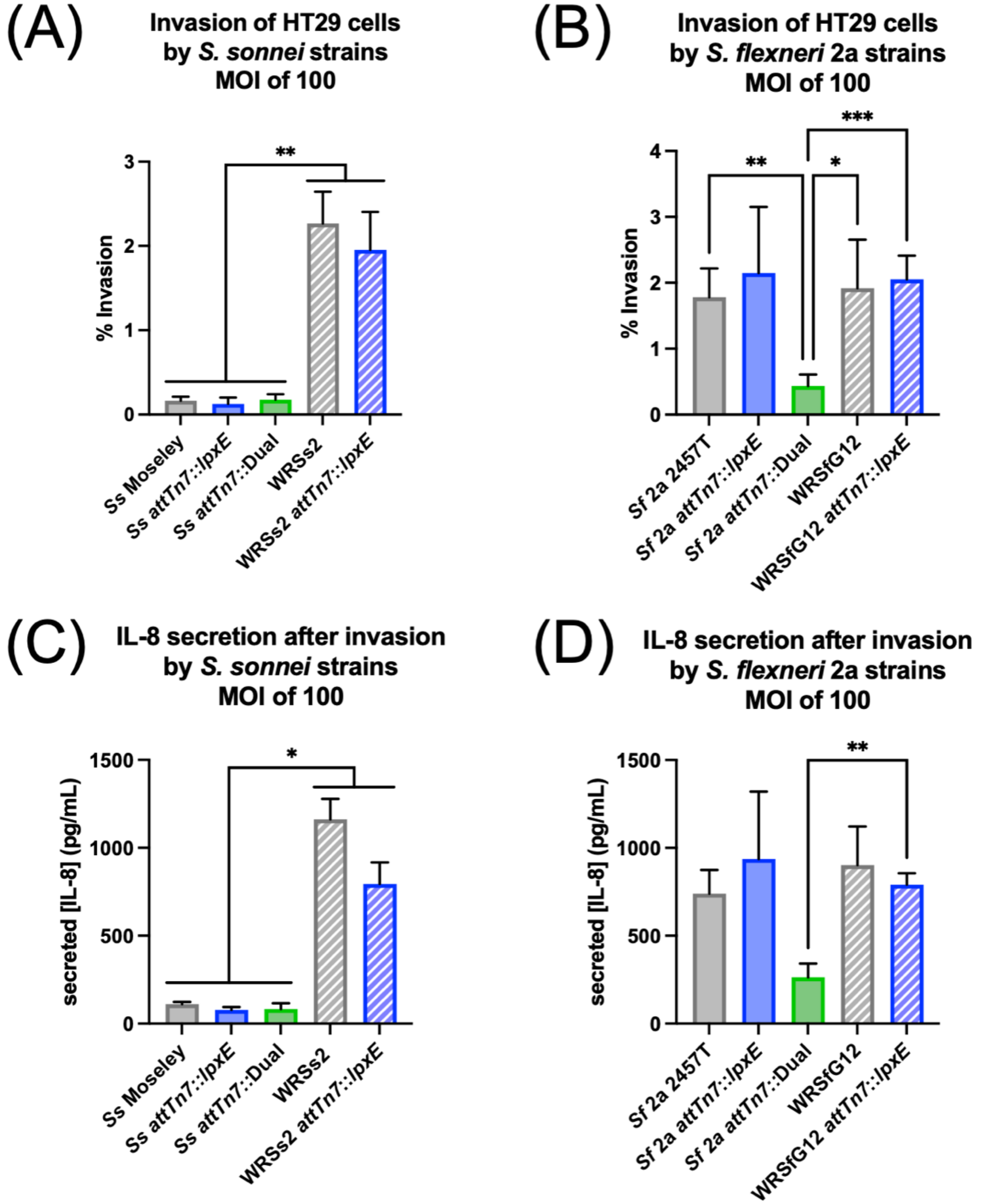

**Figure S4:** Invasion of HT29 cells by the *Shigella* strains used in this study and subsequent IL-8 secretion

(A, B) Invasion after 4 hours of exposure with *S. sonnei* and *S. flexneri* 2a unmodified and BECC-modified strains, performed on HT29 cells at an MOI 100. (C, D) IL-8 secretion in the cell supernatant after 4 hours of exposure. Statistical significance determined by ordinary one-way ANOVA. *, **, and *** represent p-values of ≤ 0.05, ≤ 0.01, and ≤ 0.001, respectively.

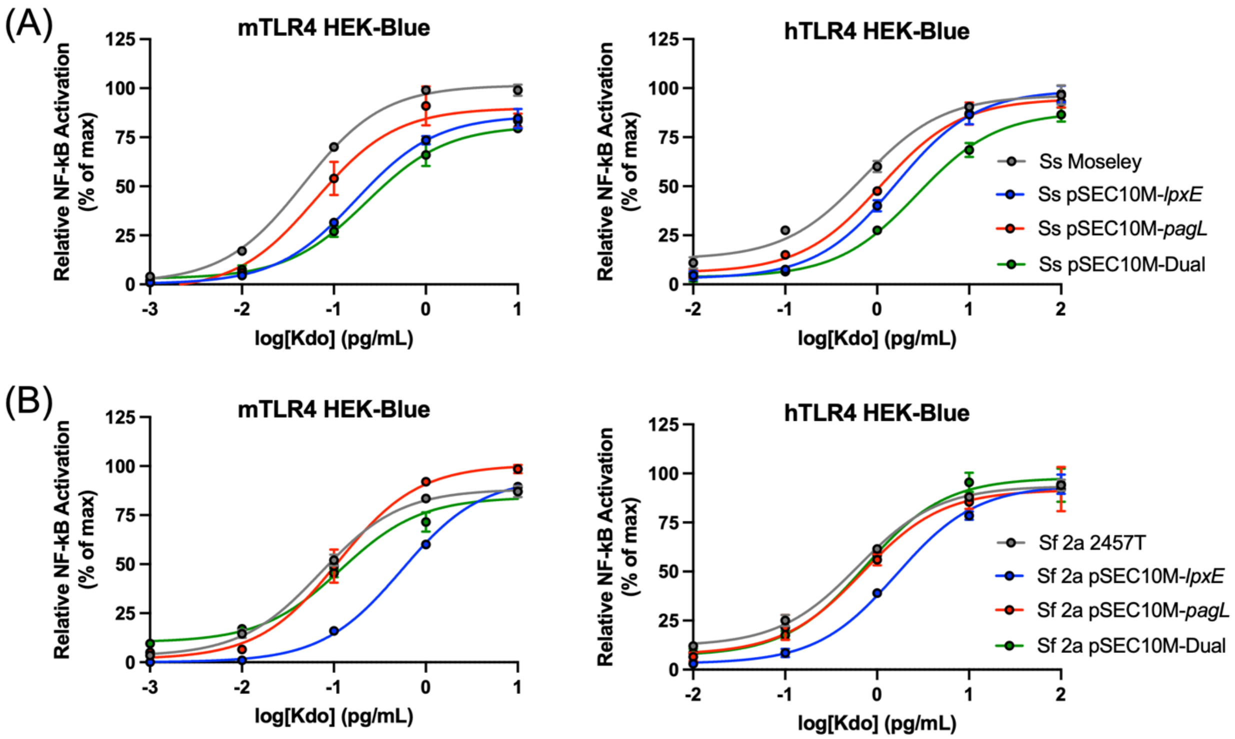

**Figure S5:** Stimulation of NF-$\kappa$B reporter cells with Kdo normalized LPS solutions from *Shigella* strains expressing BECC-enzymes from plasmids

HEK-Blue cells overexpressing human (h) and mouse (m) orthologs of TLR4/MD-2/CD-14 were stimulated across 10-fold dilutions, in duplicate, of Kdo standardized LPS for 18 hours at 37°C with 5% $CO_2$. LPS was purified from wild-type strains of (A) *S. sonnei* Moseley or (B) *S. flexneri* 2a 2457T.

**Table S1:** EC50 values for stimulation of NF-$\kappa$B reporter cells with Kdo normalized LPS solutions from *Shigella* strains expressing BECC-enzymes from plasmids

| Parent strain | Plasmid | hTLR4 HEK-Blue | | mTLR4 HEK-Blue | |
|---|---|---|---|---|---|
| | | EC50 (ng/mL) | Fold increase from unmodified | EC50 (ng/mL) | Fold increase from unmodified |
| *Ss* Moseley | - | 0.71 | - | 0.04 | - |
| | pSEC10M-*lpxE* | 1.59 | 2.23 | 0.17 | 3.72 |
| | pSEC10M-*pagL* | 1.09 | 1.54 | 0.06 | 1.36 |
| | pSEC10M-Dual | 2.71 | 3.81 | 0.22 | 4.80 |
| *Sf* 2a 2457T | - | 0.63 | - | 0.07 | - |
| | pSEC10M-*lpxE* | 1.62 | 2.59 | 0.54 | 7.61 |
| | pSEC10M-*pagL* | 0.69 | 1.11 | 0.11 | 1.55 |
| | pSEC10M-Dual | 0.73 | 1.17 | 0.11 | 1.65 |

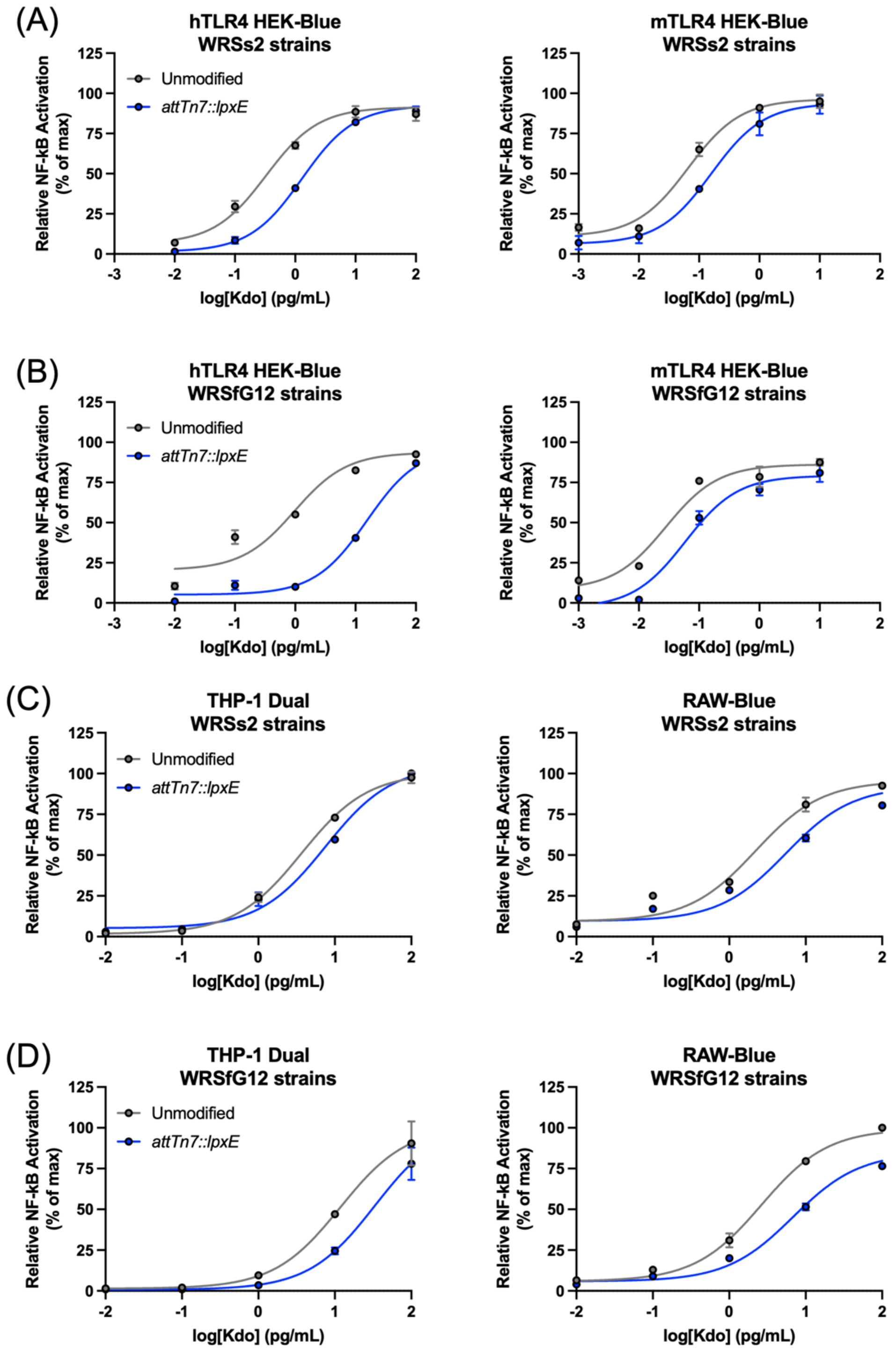

**Figure S6:** Stimulation of NF-$\kappa$B reporter cells with LPS solutions from *attTn7* integrants of the vaccine strains of *Shigella* used in this study.

Human- and mouse-HEK-Blue, RAW-Blue, and THP-1 Dual cells were stimulated across 10-fold dilutions, in duplicate, of Kdo standardized LPS for 18 hours at 37°C with 5% $CO_2$ with purified LPS from vaccine strains (A, C) WRSs2 and (B, D) WRSfG12.

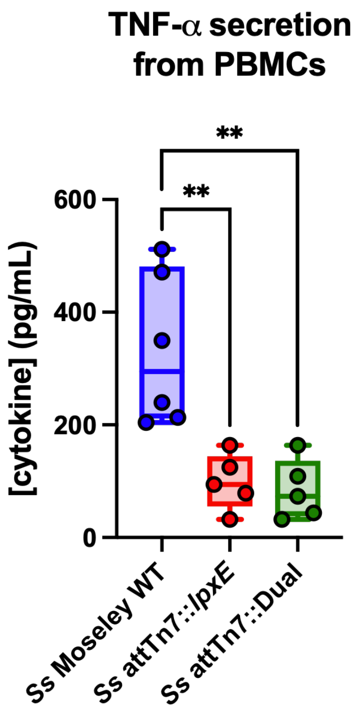

**Figure S7:** PBMC stimulation with purified LPS samples from wild-type *S. sonnei* Moseley

Human PBMCs from 6 independent donors were stimulated at 1 pg/mL Kdo for 18 hours at 37°C with 5% $CO_2$ with purified LPS from wild-type strains of *S. sonnei* Moseley and TNF-α secretion in the supernatant quantified by cytokine ELISA. Statistical significance determined by ordinary one-way ANOVA. ** represents a p-value ≤ 0.01.

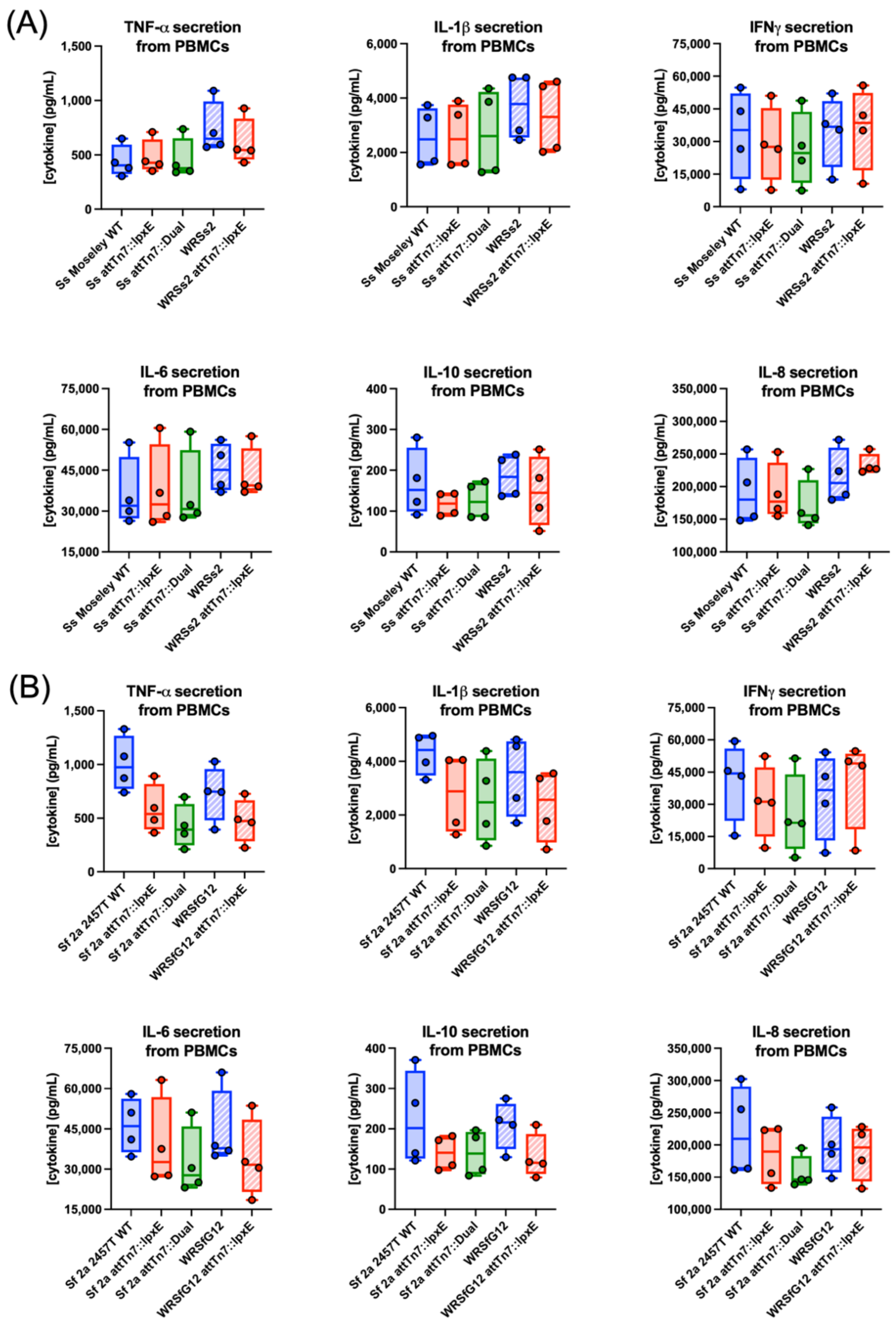

**Figure S8:** PBMC stimulation with purified LPS from both wild-type and attenuated *S. sonnei* and *S. flexneri* 2a strains used in this study.

Human PBMCs from 4 independent donors were stimulated at 1 pg/mL Kdo for 48 hours at 37°C with 5% $CO_2$ with purified LPS from wild-type and attenuated strains of (A) *S. sonnei* Moseley and (B) *S. flexneri* 2a 2457T. Supernatant cytokine levels were quantified by multiplex analysis. The top six cytokines produced upon LPS stimulation are displayed as box and whisker plots.

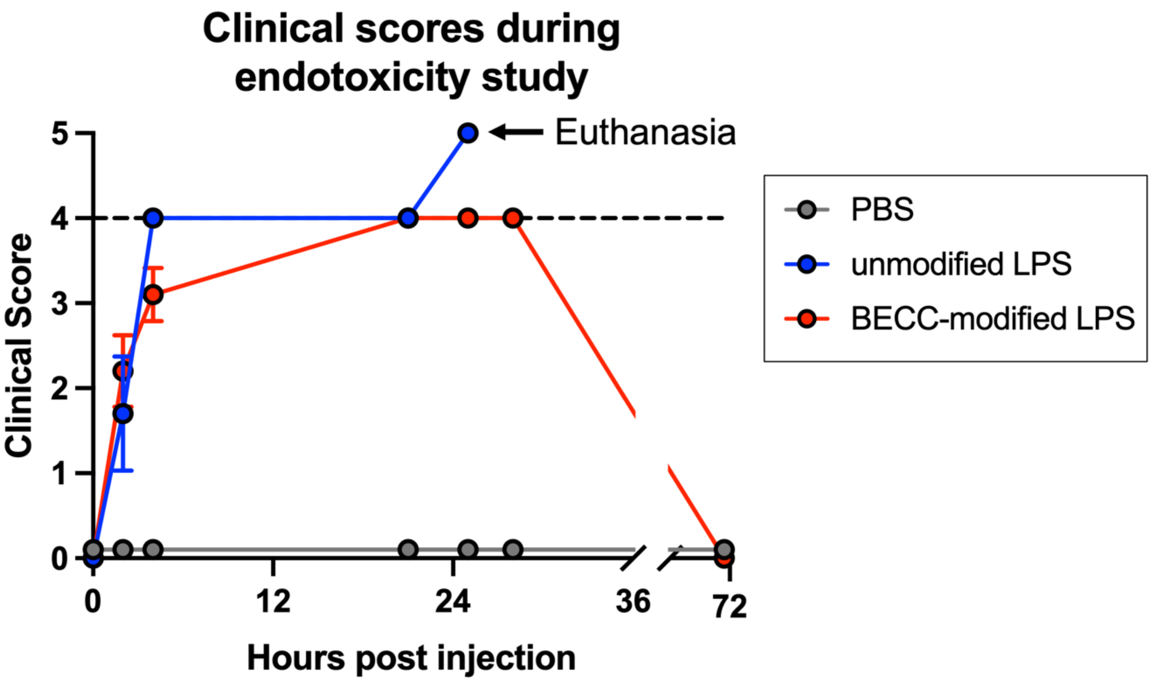

**Figure S9:** Clinical scores throughout acute endotoxemia model assessing endotoxicity of LPS

A representative plot of clinical scores for a group of 10 mice who received intraperitoneal injections of a $Kdo_2$ normalized dose of LPS from WRSs2, representative of 15 mg/kg. Average clinical scores ± SD throughout the study, whereby mice were euthanized once they reached a clinical score of 5.

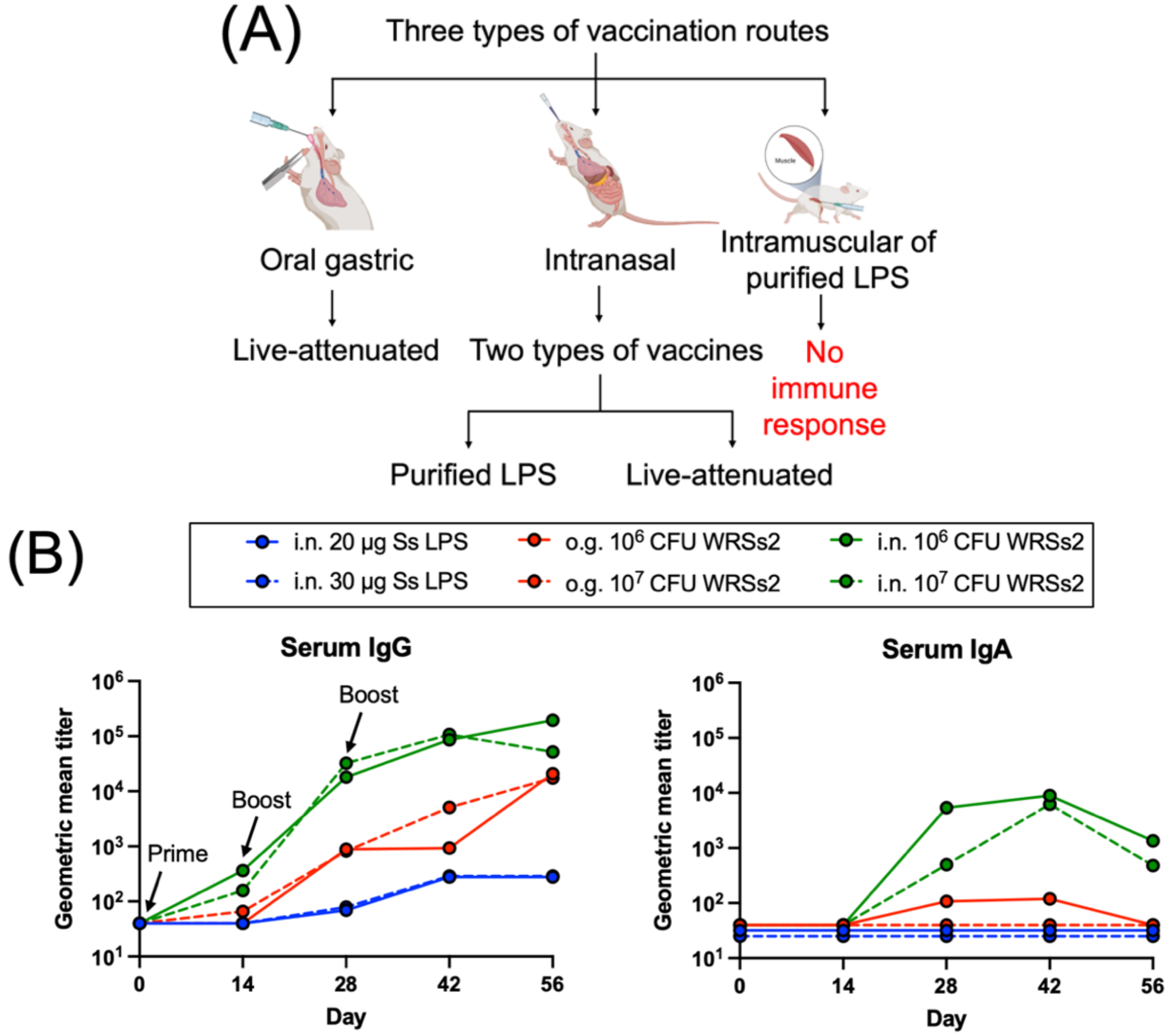

**Figure S10:** Optimization of the *Shigella* murine vaccination model

(A) Various immunization routes were tested using either WRSs2 or purified LPS from wild-type *S. sonnei* Moseley. (B) Serum IgG and IgA antibody titers over time against wild-type *S. sonnei* Moseley LPS. Higher dose groups are indicated by a dashed line. Arrows indicate administration of vaccine doses. Errors bars are omitted for clarity.

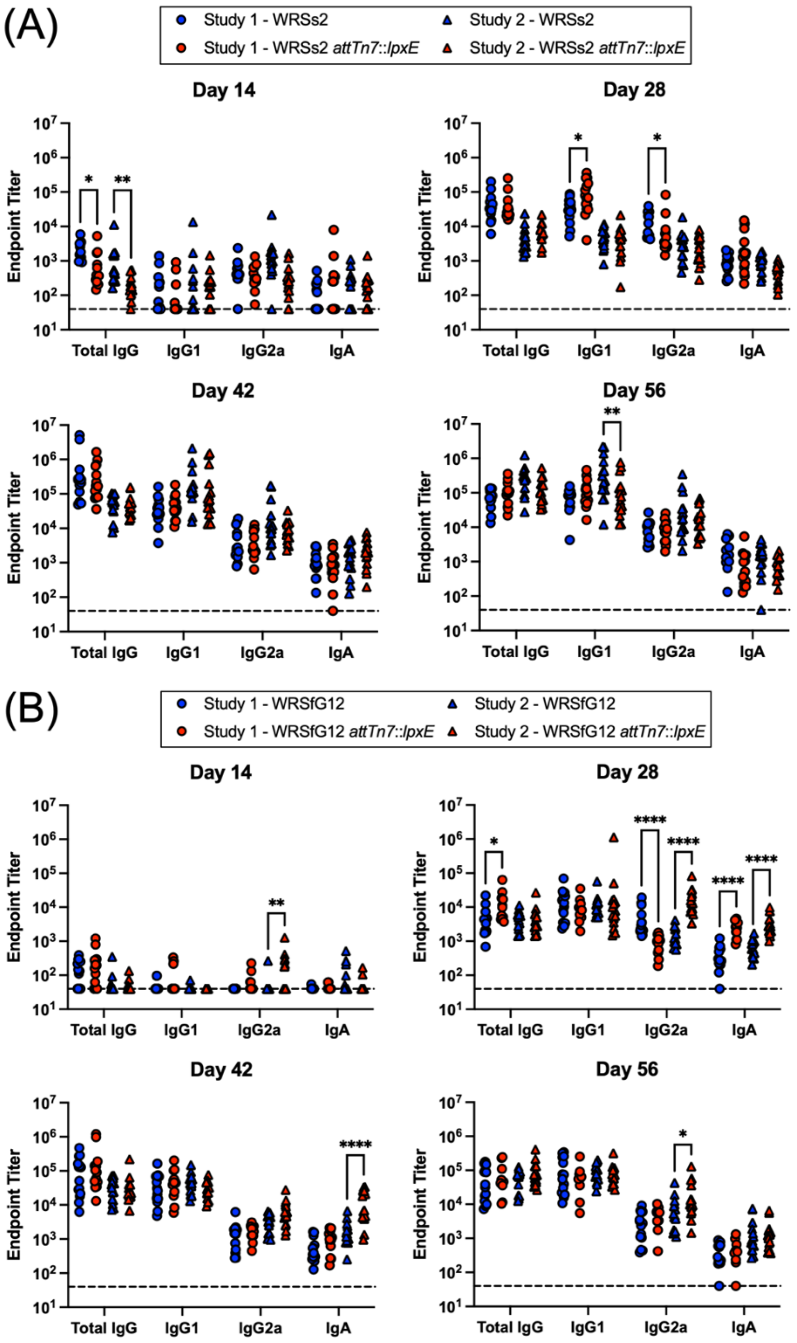

**Figure S11:** Antibody titers from two-independent *Shigella* murine vaccine studies

Serum IgG, IgG1, IgG2a, and IgA antibody titers for each group (n=15) against (A) *S. sonnei* Moseley or (B) *S. flexneri* 2a 2457T LPS. Antibody titers for each individual mouse at all time points of the vaccine study are shown. Circles and triangles, indicate titers from the first and second study, respectively. The dashed line indicates a value where no response was observed at the lowest dilution tested for the ELISA. Statistical significance determined by 2way ANOVA. *, **, ***, **** represent p-values of ≤ 0.05, ≤ 0.01, ≤ 0.001, and ≤ 0.0001, respectively.

**Table S2:** Total IgG- and IgA-ASCs per $10^6$ spleen/lung cells induced upon LPS stimulation

| Vaccine | Study 1 | | | | Study 2 | | | |
|---|---|---|---|---|---|---|---|---|
| | ASCs in spleens | | ASCs in lungs | | ASCs in spleens | | ASCs in lungs | |
| | Mean[c] IgG (range) | Mean IgA (range) | Mean IgG (range) | Mean IgA (range) | Mean IgG (range) | Mean IgA (range) | Mean IgG (range) | Mean IgA (range) |
| PBS[a] | 169 (150-190) | 49 (30-80) | 14 (10-20) | 10 (10-10) | 80 (0-150) | 40 (0-100) | 50 (0-100) | 90 (50-150) |
| WRss2[a] | 399 (340-500) | 88 (60-130) | 71 (30-110) | 75 (10-190) | 130 (20-600) | 65 (20-160) | 32 (0-100) | 105 (80-150) |
| WRSs2 *attTn7*::*lpxE*[a] | 246 (24-580) | 49 (8-270) | 35 (10-70) | 35 (10-240) | 298 (60-1250) | 90 (40-160) | 52 (30-100) | 140 (70-350) |
| PBS[b] | 317 (0-500) | 46 (0-100) | 10 (0-20) | 57 (20-80) | 70 (0-150) | 40 (0-100) | 30 (0-150 | 90 (50-100) |
| WRSfG12[b] | 360 (130-600) | 81 (50-120) | 98 (60-140) | 68 (40-210) | 122 (20-500) | 50 (30-140) | 92 (0-200) | 112 (30-200) |
| WRSfG12 *attTn7*::*lpxE*[b] | 391 (120-600) | 65 (30-120) | 55 (20-140) | 88 (60-150) | 152 (60-650) | 64 (10-350) | 53 (20-100) | 91 (50-200) |

[a]Wild-type *S. sonnei* Moseley LPS used as antigen for stimulation
[b]Wild-type *S. flexneri* 2a 2457T LPS used as antigen for stimulation
[c]Means calculated as geometric means

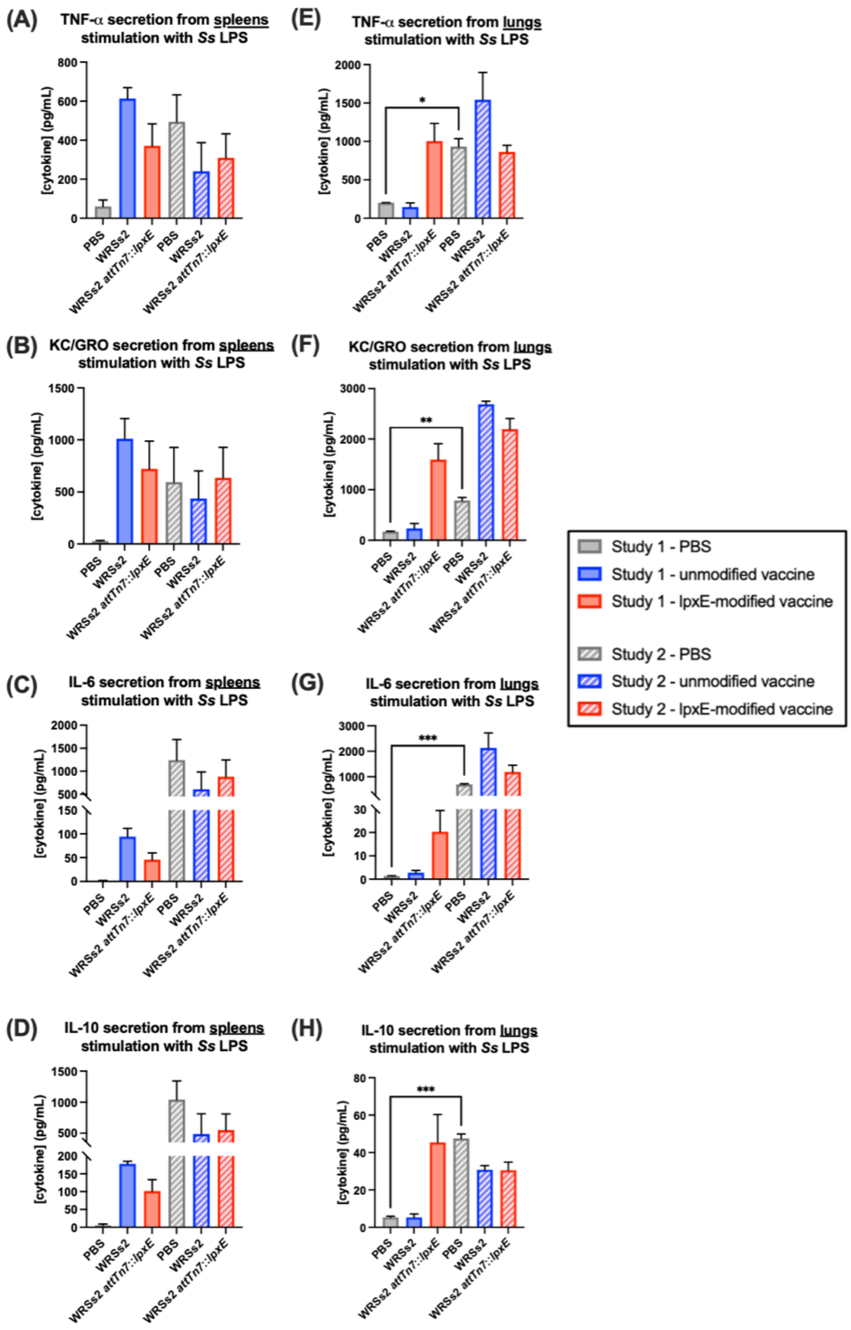

**Figure S12:** LPS-specific cytokine responses from spleens and lungs of mice vaccinated with parental and LpxE-modified WRSs2

(A-D) Spleens and (E-H) lungs were obtained on day 56 from 5 mice in the indicated groups and single-cell suspensions were prepared. Cells were incubated for 48 h with *S. sonnei* Moseley wild-type LPS, and supernatant cytokine levels measured by multiplex analysis. Of all cytokines measured, the four with the greatest abundance are shown as the mean concentrations +SEM. Study one is represented by filled boxes and study two by striped boxes. Statistical significance determined by One-way ANOVA with Brown-Forsythe and Welch corrections. *, **, and *** represent p-values ≤ 0.05, ≤ 0.01, and ≤ 0.001, respectively.

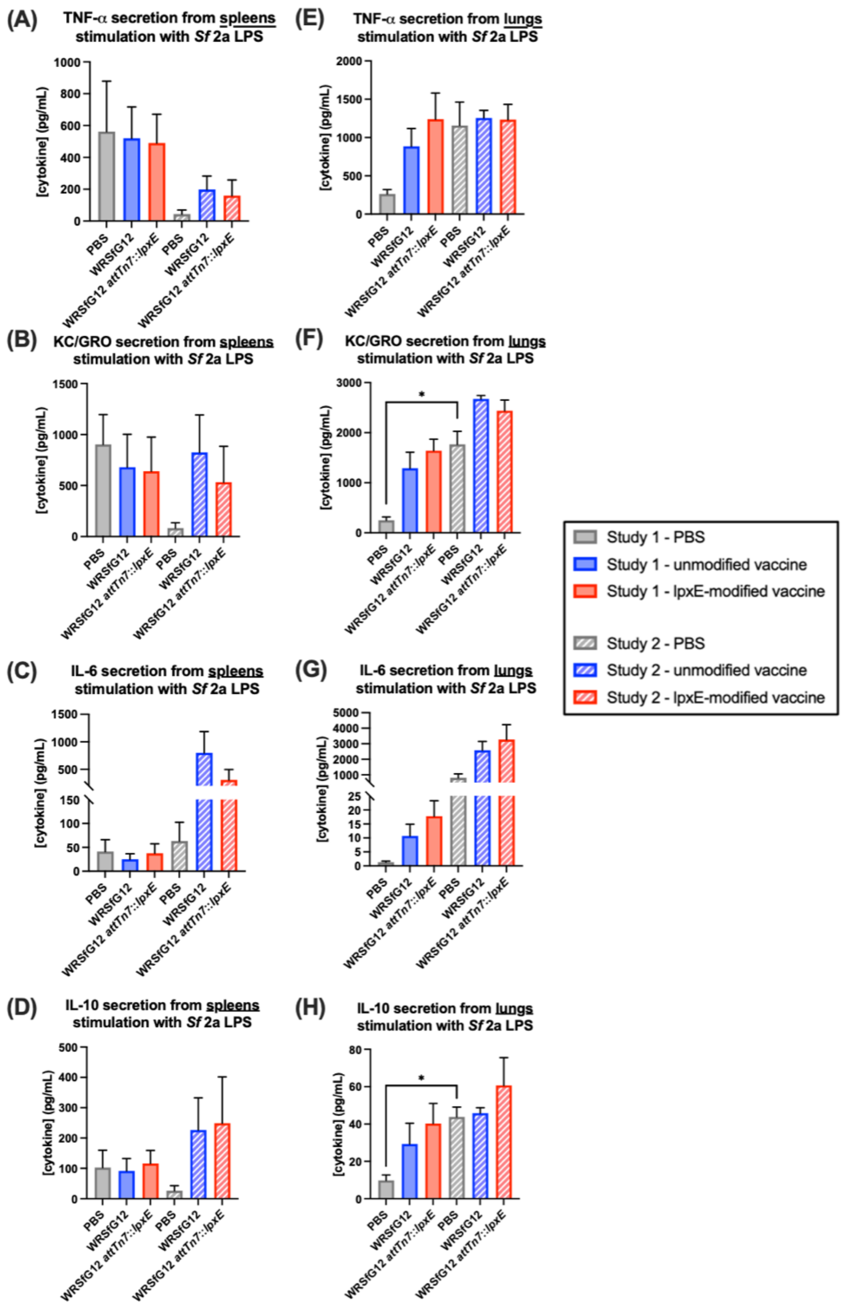

**Figure S13:** LPS-specific cytokine responses from spleens and lungs of mice vaccinated with parental and LpxE-modified WRSfG12

(A-D) Spleens and (E-H) lungs were obtained on day 56 from 5 mice in the indicated groups and single-cell suspensions were prepared. Cells were incubated for 48 h with *S. flexneri* 2a 2457T wild-type LPS, and supernatant cytokine levels measured by multiplex analysis. Of all cytokines measured, the four with the greatest abundance are shown as the mean concentrations +SEM. Study one is represented by filled boxes and study two by striped boxes. Statistical significance determined by One-way ANOVA with Brown-Forsythe and Welch corrections. *, **, and *** represent p-values ≤ 0.05, ≤ 0.01, and ≤ 0.001, respectively.

**Table S3:** Bacterial strains used in this study

| Strain | Description | Genotype | Modified lipid A | Ref. |
|---|---|---|---|---|
| *S. sonnei* Moseley | *S. sonnei* parent strain | Wild-type | - | [1] |
| *S. flexneri* 2a 2457T | *S. flexneri* 2a parent strain | Wild-type | - | [2] |
| WRSs1 | 1st generation attenuated vaccine strain of *S. sonnei* | *ΔvirG* | - | [3] |
| WRSs2 | 2nd generation attenuated vaccine strain of *S. sonnei* | *ΔvirG, senA, senB* | - | [1,4-8] |
| SC602 | 1st generation attenuated vaccine strain of *S. flexneri* 2a | *ΔvirG, iuc* | - | [9,10] |
| WRSfG12 | 2nd generation attenuated vaccine strain of *S. flexneri* 2a | *ΔvirG, iuc, setAB, senA, senB* | - | [6,11] |
| *E. coli* S17-1 | Mobilizes *oriT*-carrying plasmids for conjugal transfer | *recA, pro, RP4-2 Tet::Mu-Kan::Tn7* | - | [12] |
| *E. coli* S17-1 pGRG36-*lpxE* | For conjugation of pGRG36-*lpxE* | pGRG36::$P_{ompC}$–*lpxE* | - | This study |
| *E. coli* S17-1 pGRG36-Dual | For conjugation of pGRG36-Dual | pGRG36::$P_{ompC}$–Dual | - | This study |
| *E. coli* DH5α | For plasmid propagation | *F⁻ Φ80lacZΔM15 Δ(lacZYA-argF)U169 recA1 endA1 hsdR17 ($r_k^-$, $m_k^+$) phoA supE44 thi-1 gyrA96 relA1 λ⁻* | - | Thermo Fisher |
| *E. coli* TOP10 | For plasmid propagation | *F- mcrA Δ(mrr-hsdRMS-mcrBC) Φ80lacZΔM15 ΔlacX74 recA1 araD139 Δ(araleu)7697 galU galK rpsL (StrR) endA1 nupG* | - | Thermo Fisher |
| *S. sonnei* pSEC10M-*pagL* | Wild-type strain, plasmid-based expression of *pagL* under control of $P_{ompC}$ | pSEC10*M*::$P_{ompC}$–*pagL* | + | This study |
| *S. sonnei* pSEC10M-*lpxE* | Wild-type strain, plasmid-based expression of *lpxE* under control of $P_{ompC}$ | pSEC10M::$P_{ompC}$–*lpxE* | + | This study |
| *S. sonnei* pSEC10M-Dual | Wild-type strain, plasmid-based expression of Dual under control of $P_{ompC}$ | pSEC10M::$P_{ompC}$–*lpxE–pagL* | + | This study |
| *S. flexneri* 2a pSEC10M-*pagL* | Wild-type strain, plasmid-based expression of *pagL* under control of $P_{ompC}$ | pSEC10M::$P_{ompC}$–*pagL* | + | This study |
| *S. flexneri* 2a pSEC10M-*lpxE* | Wild-type strain, plasmid-based expression of *lpxE* under control of $P_{ompC}$ | pSEC10M::$P_{ompC}$–*lpxE* | + | This study |

| Strain | Description | Genotype | Modified lipid A | Ref. |
|---|---|---|---|---|
| *S. flexneri* 2a pSEC10M-Dual | Wild-type strain, plasmid-based expression of Dual under control of $P_{ompC}$ | pSEC10M::$P_{ompC}$–*lpxE*–*pagL* | + | This study |
| WRSs1 pSEC10M-*pagL* | Attenuated 1st generation vaccine strain of *S. sonnei*, plasmid-based expression of *pagL* under control of $P_{ompC}$ | Δ*virG;* pSEC10M::$P_{ompC}$–*pagL* | + | This study |
| WRSs1 pSEC10M-*lpxE* | Attenuated 1st generation vaccine strain of *S. sonnei*, plasmid-based expression of *lpxE* under control of $P_{ompC}$ | Δ*virG;* pSEC10M::$P_{ompC}$–*lpxE* | + | This study |
| SC602 pSEC10M-*pagL* | Attenuated 1st generation vaccine strain of *S. flexneri* 2a, plasmid-based expression of *pagL* under control of $P_{ompC}$ | Δ*virG, iuc;* pSEC10M::$P_{ompC}$–*pagL* | + | This study |
| SC602 pSEC10M-*lpxE* | Attenuated 1st generation vaccine strain of *S. flexneri* 2a, plasmid-based expression of *lpxE* under control of $P_{ompC}$ | Δ*virG, iuc;* pSEC10M::$P_{ompC}$–*lpxE* | + | This study |
| *S. sonnei attTn7*::$P_{ompC}$–*lpxE* | Wild-type strain, chromosomal expression of *lpxE* under control of $P_{ompC}$ from *attTn7* site from *attTn7* site | *attTn7*::$P_{ompC}$–*lpxE* | + | This study |
| *S. sonnei attTn7*::$P_{ompC}$–Dual | Wild-type strain, chromosomal expression of Dual under control of $P_{ompC}$ from *attTn7* site | *attTn7*::$P_{ompC}$–*lpxE*–*pagL* | + | This study |
| *S. flexneri* 2a *attTn7*::$P_{ompC}$–*lpxE* | Wild-type strain, chromosomal expression of *lpxE* under control of $P_{ompC}$ from *attTn7* site | *attTn7*::$P_{ompC}$–*lpxE* | + | This study |
| *S. flexneri* 2a *attTn7*::$P_{ompC}$–Dual | Wild-type strain, chromosomal expression of Dual under control of $P_{ompC}$ from *attTn7* site | *attTn7*::$P_{ompC}$–*lpxE*–*pagL* | + | This study |
| WRSs2 *attTn7*::$P_{ompC}$–*lpxE* | Attenuated 2nd generation vaccine strain of *S. sonnei*, chromosomal based expression of *lpxE* under control of $P_{ompC}$ from *attTn7* site | Δ*virG, senA, senB; attTn7*::$P_{ompC}$–*lpxE* | + | This study |
| WRSfG12 *attTn7*::$P_{ompC}$–*lpxE* | Attenuated 2nd generation vaccine strain of *S. flexneri* 2a, chromosomal based expression of *lpxE* under control of $P_{ompC}$ from *attTn7* site | Δ*virG, iuc, setAB, senA, senB; attTn7*::$P_{ompC}$–*lpxE* | + | This study |

**Table S4:** Primers used in this study

| Primer | Sequence 5' → 3' | Reference |
|---|---|---|
| *ospD3*-F<br>*ospD3*-R | TACACGTCCATTATGCAAGGCT<br>TGCCATCAGTAAATTTAATCCCATC | [13] |
| *ipaB*-F<br>*ipaB*-R | CACAGCATCTGCTGAACAGC<br>CAGCAGAAGCGACACTTCCT | This study |
| *lpxE*-F<br>*lpxE*-R | TGCCATCAGTAAATTTAATCCCATC<br>GGGGGCTAGCTTAAATAATCTCTCTATTTCTCATCT | This study |
| *pagL*-F<br>*pagL*-R | GGGGGAATTCGGATCCGTGTATATGAAGAG<br>GGGGGCTAGCTCAGAAATTATAACTAATTG | This study |
| $P_{ompC}$-F | GAATTCTGTGGTAGCACAGA | This study |
| pSEC10M-F<br>pSEC10M-R | TTTTGAATTCGCGGCCGCATTTAAATGCTAGCAAAA<br>TTTTGCTAGCATTTAAATGCGGCCGCGAATTCAAAA | This study |
| RT-qPCR *lpxE*-F<br>RT-qPCR *lpxE*-R | TTCCATCACCGTTGGAGCAA<br>TAGCAACCAAAAGCAACGCC | This study |
| RT-qPCR *rpoA*-F<br>RT-qPCR *rpoA*-R | CGTATCAAAGTTCAGCGCGG<br>TCGATGACCAGCTTGTCCAG | This study |

**Table S5:** Plasmids used in this study

| Plasmid | Description | Genotype | Ref. |
|---|---|---|---|
| pSEC10 | Osmotic-inducible vector for gene expression under $P_{ompC}$ | *ori101*, *hok-sok*, *parA/B;* $P_{ompC}$–*clyA* | [14] |
| pSEC10M | Modified pSEC10 plasmid with $P_{ompC}$ and *clyA* replaced by an MCS | *ori101*, *hok-sok*, *parA/B*; $P_{ompC}$–*clyA*::MCS | This study |
| pGRG25 | Arabinose-inducible Tn7 transposition vector with MCS site | *ori ts, oriT, araC,* $P_{BAD}$–*tnsABCD*, *mTn7*::MCS | [15] |
| pGRG36 | Arabinose-inducible Tn7 transposition vector with SmaI added to MCS site of pGRG25 | *ori ts, oriT, araC,* $P_{BAD}$–*tnsABCD*, *mTn7*::MCS | This study |

**Table S6:** Clinical scores for mice during endotoxicity study

| Score | Description | Appearance | Mobility |
|---|---|---|---|
| 0 | Perfectly healthy | - | - |
| 1 | Smooth | Slightly ruffled coat | Active and scurrying, burrowing |
| 2 | Slightly ruffled | Ruffled coat | Active and scurrying, burrowing |
| 3 | Ruffled | Very ruffled coat | Walking, but no scurrying, mildly lethargic |
| 4 | Sick | Very ruffled coat, inset eyes | Slow to no movement, extremely lethargic |
| 5 | Very sick (euthanize) | Very ruffled coat, closed inset eyes | No movement or spastic movements, will not return upright if put on side, noticeable stress |
| 6 | Deceased | - | - |
